# Structural Characterization of the Type IV Secretion System in *Brucella melitensis* for Virtual Screening-Based Therapeutic Targeting

**DOI:** 10.64898/2026.02.18.706537

**Authors:** Jahnvi Kapoor, Amisha Panda, Raman Rajagopal, Sanjiv Kumar, Anannya Bandyopadhyay

**Affiliations:** Protein Biology Lab, Department of Zoology, University of Delhi, Delhi, India; Gut Biology Laboratory, Department of Zoology, University of Delhi, New Delhi, India; Independent researcher, Stockholm, Sweden

**Keywords:** *Brucella melitensis*, Brucellosis, Type IV Secretion System, VirB proteins, Structural Modeling, Protein-Protein interactions

## Abstract

Brucellosis is a globally important zoonotic disease caused by *Brucella melitensis*, the most virulent and clinically significant species affecting both humans and livestock. Unlike many Gram-negative pathogens, *B. melitensis*, a facultative intracellular pathogen, lacks conventional virulence factors and instead relies on specialized systems such as the Type IV Secretion System (T4SS) for secretion of effector proteins. In this study, an integrated computational pipeline was implemented to identify, model, and assemble the T4SS components, encoded by *virB* operon, from the complete *B. melitensis* proteome. Template-based modeling strategies were employed to generate structures of T4SS subcomplexes, referencing crystallographic data from *E. coli* T4SS. Structural superposition with *E. coli* homologs revealed highly conserved architecture despite only 30–50% sequence identity. Stereochemical validation confirmed high model quality and favorable interactions among most VirB protein pairs. Membrane insertion analysis of the membrane-embedded assemblies further corroborated the spatial orientation of the modeled T4SS. Potential of T4SS as a drug target was explored by targeting dimeric interface of VirB11 ATPase to disrupt protein-protein interactions that could disarm the pathogen. Virtual screening of compounds from DrugBank database revealed compounds with docking score ≤ -7.0 kcal/mol that were screened based on ADMET properties, yielding three promising candidates – Ezetimibe (Drug Id: DB00973), Chlordiazepoxide (Drug Id: DB00475), and Alloin (Drug Id: DB15477). MM-GBSA analysis estimated favorable binding free energies for these compounds and molecular dynamics simulation for 200 ns further confirmed the protein-ligand interaction stability. Collectively, these findings provide new insights into the architecture of *B. melitensis* T4SS and identify three potential drug molecules targeting T4SS. This supports FDA – approved drug repurposing as an effective strategy for anti-virulence therapy against Brucellosis.

**Graphical Abstract:** 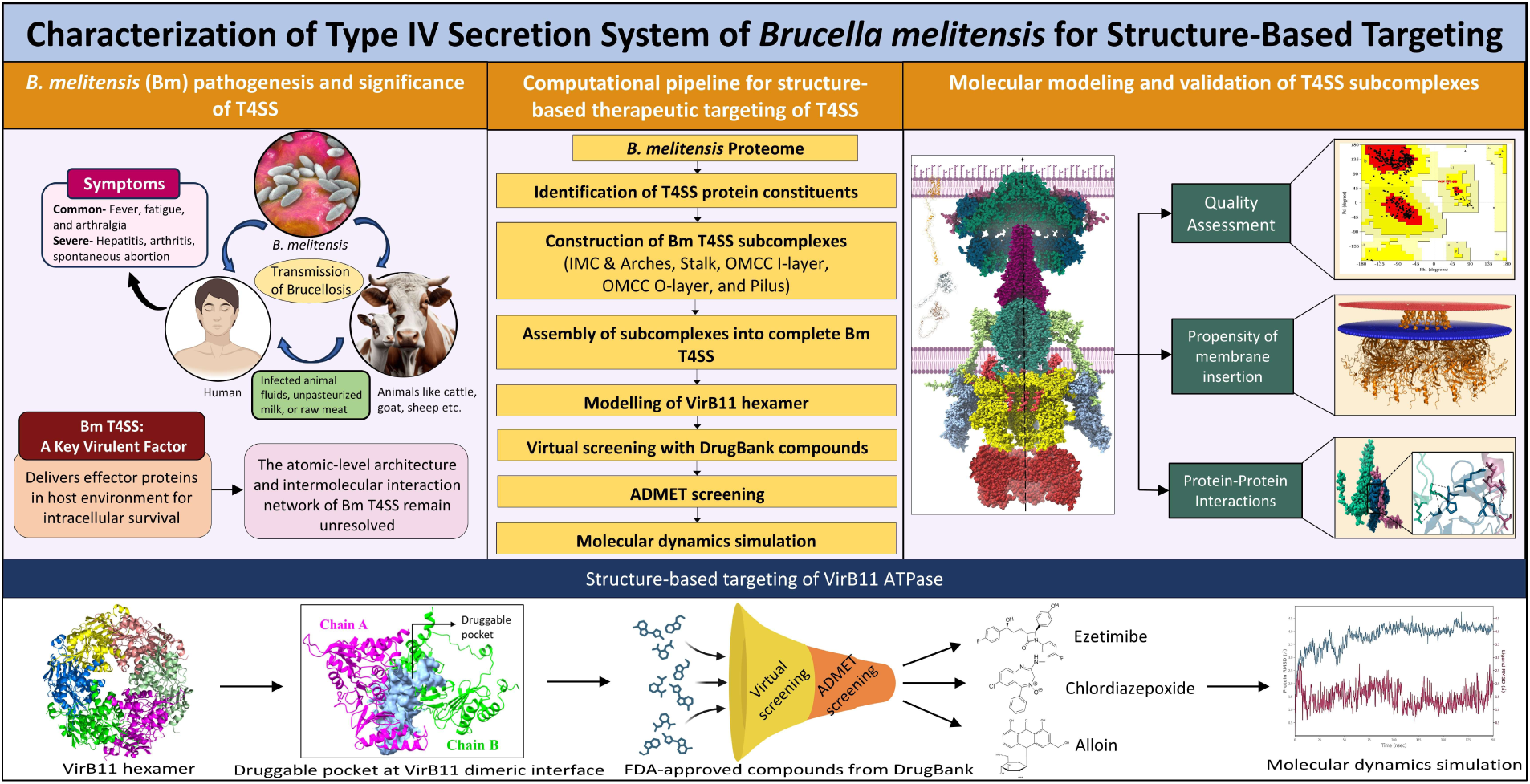

## 1. Introduction

Brucellosis (Malta fever) is a zoonotic infection caused by various *Brucella* spp. that afflicts both livestock and humans. The disease affects 1.6–2.1 million people globally each year, particularly in Africa, Asia, and the Mediterranean region [1,2]. Clinical symptoms range from fever, fatigue, and joint pain to severe complications like arthritis, hepatitis, and pregnancy-related issues [1,3]. *Brucella* are small (0.5–0.7 μm × 0.6–1.5 μm), aerobic, Gram-negative, non-motile coccobacilli [4] with an incubation period ranging from 5 days to 6 months [5]. *Brucella melitensis* (Bm) is a facultative intracellular pathogen and the most virulent species causing brucellosis in humans [3]. Unlike many Gram-negative pathogens, Bm lacks conventional virulence determinants such as exotoxins, cytolysins, exoenzymes, plasmids, fimbriae, and drug-resistant forms. Instead, its pathogenicity primarily depends on key virulence systems including lipopolysaccharide (LPS), the Type IV Secretion System (T4SS), and the BvrR/BvrS two-component regulatory system [6]. These systems collectively mediate host-pathogen interaction, and the establishment of *Brucella*-containing vacuoles (BCVs) where bacterial replication occurs [6].

Bacterial secretion systems are often described as molecular “nanomachines” essential for establishing infection, and facilitating the delivery of virulence factors. The general secretion (Sec) and twin arginine translocation (Tat) pathways are universal bacterial secretion systems essential for protein export across the cytoplasmic membrane [7]. Beyond the Sec and Tat pathways, bacteria have evolved other specialized secretion systems, classified as types I through XI, that mediate the secretion of effector proteins across the outer membrane or directly into the target cells [8]. The Type IV Secretion System (T4SS) is an important virulence factor in several bacterial pathogens, such as *Helicobacter pylori*, *Legionella pneumophila*, *Bartonella* spp. and *Brucella* spp. [9]. The T4SS in *Brucella* spp. has been shown to be an essential component, as mutants of *B. abortus*, *B. suis*, and *B. melitensis* lacking a functional T4SS exhibit marked attenuation in both intracellular survival assays and mouse infection models [10–16]. Bm T4SS secretes effector proteins, such as RicA, BtpA, VceA, VceC, BspA, BspB, and BspF, which perform diverse biological functions for intracellular survival [17,18]. The Bm T4SS is encoded by the *virB* operon on chromosome II where *virB1* to *virB12* form a transcriptional unit that encodes for 12 VirB family proteins out of which eight have been annotated as VirB in NCBI database [10,19]. The architectures of T4SS from two closely related bacterial species, one from *Agrobacterium tumefaciens* (VirB/D4 system) and another from *E. coli* (encoded on the R388 conjugative plasmid), are well established. The overall structure of T4SS is composed of five subcomplexes: Inner membrane complex (IMC) and arches (VirB3, VirB4, and VirB8); the stalk (VirB5, and VirB6); outer membrane core complex inner layer (OMCC I-layer: VirB9, and VirB10); outer membrane core complex outer layer (OMCC O-layer: VirB7, VirB9, and VirB10); and the pilus (VirB2) [20,21]. In *E. coli* (Ec) T4SS, the IMC and arches subcomplex assembles into a hexameric scaffold embedded in the inner membrane. The IMC is composed of a central ATPase that energizes the translocation of effector proteins across the inner membrane. The stalk forms a pentameric subcomplex that creates a continuous channel connecting IMC to the OMCC subcomplex. In the outer membrane, the OMCC presents a barrel shaped architecture which is divided into two layers that are asymmetric to each other enabling the unidirectional translocation of substrate molecules [20,22]. Further, the pilus consists of a polymer of VirB2 homolog that forms a cylindrical structure on the bacterial surface [23]. Studies have shown that VirB2 proteins assemble at VirB5-VirB6 interface in stalk subcomplex, and push the VirB5 pentamer upwards to open the initially closed OMCC gate [18,24,25]. Even though the complete structure of Bm T4SS has not been characterized like that of Ec, studies have shown some similarities and differences between the two species. The Bm T4SS is a 94-subunit macromolecular assembly that relies on extensive protein–protein interactions (PPI) for its proper functioning. In Bm T4SS, the protomer of IMC and arches subcomplex have two VirB8 subunits, whereas Ec T4SS features three copies of VirB8 homolog [21,26]. The IMC is composed of a central VirB4 ATPase which is anchored to VirB11 ATPase on the cytoplasmic side [27]. VirB11 is an essential cytoplasmic ATPase that act as a “switch” to provide energy for secretion of effector proteins. VirB11 has been crystallized in *B. suis* (PDB Id: 2GZA) [27]. A previous study in *B. abortus* had shown that attenuation of VirB11 leads to defective trafficking of vesicles to ER, resulting in impaired bacterial replication within macrophages, and consequent avirulence in mice models [28]. PPI can be targeted to inhibit VirB11 assembly that could disarm the pathogen and the infection can eventually be cleared by the host [26,29,30]. Further, in periplasm, the stalk subcomplex consists of pentamers of each VirB5 and VirB6 forming a bridge between the IMC, and the OMCC subcomplexes [31]. Notably, the OMCC architecture in Bm T4SS remains poorly characterized. Other components of Bm T4SS include VirB1 and VirB12. VirB1 is a lytic transglycosylase that does not associate with any other T4SS protein [23]. In *A. tumefaciens*, VirB1 is involved in destruction of peptidoglycan layer in periplasm to facilitate the T4SS assembly [32]. VirB12 is a surface-exposed protein in *Brucella* spp. that contains an OmpA-like homology domain [33]. Although its precise function remain unclear, it is thought to play a role in mediating adhesion and interaction with host cells [33].

Existing studies in *Brucella* spp. have focused on individual VirB proteins for vaccine development and serological diagnosis [34–37] but the complete architecture of Bm T4SS has not yet been determined. In this study, we have applied a computational pipeline to identify and characterize all the constituent proteins of Bm T4SS. VirB protein sequences were identified from the complete proteome of Bm using three approaches: Operon-based genomic localization, TXSScan tool, and sequence similarity with VirB homologs in other *Brucella* spp. The monomeric structures of VirB proteins were modeled using AlphaFold 3. The constituent proteins were assembled into subcomplexes based on structural homology with Ec T4SS architecture. The binding affinity and intermolecular protein interactions were determined for each subcomplex. These interactions served as promising targets for the development of therapeutic inhibitors aimed at disrupting the activity of the secretion system. Amino acid residues forming a druggable pocket were identified at VirB11 dimeric interface. Structure-based virtual screening was performed against identified pocket using compounds from DrugBank database. Selected compounds were then filtered based on their absorption, distribution, metabolism, excretion, and toxicity (ADMET) properties. Energetics and stability of the VirB11 dimer – ligand complexes were assessed using MM-GBSA method and molecular dynamics simulation. Overall, the study identified three promising candidates with the potential to attenuate pathogen virulence and aid in the control of Brucellosis.

## 2. Materials and Methods

### 2.1 Identification of T4SS constituent proteins

The reference strain *B. melitensis* biovar 1 strain 16M was chosen for this study. Amino acid sequences of all the proteins of Bm 16M were downloaded from NCBI [19] using genome assembly ASM74041v1 (accessed on 25 August 2025).

To identify all the proteins of the Bm T4SS, three complementary strategies were employed. The computational pipeline is detailed below and schematically represented in Figure 1. Initially, annotated VirB proteins were identified based on genomic location data from the operon available in the NCBI database. Since not all the proteins were annotated as VirB family proteins, other approaches were employed. In the second approach, TXSScan [38] tool from MacSyFinder was used for annotation of bacterial secretion system proteins (accessed on 28 August 2025). FASTA file of complete proteome was submitted to the Galaxy server that hosts TXSScan tool (https://galaxy.pasteur.fr/). Furthermore, remaining VirB proteins were identified based on sequence similarity with their homologs present in *B. abortus* (accession no.-AAF73894.1 and AAF73905.1) [33] using BLASTP.

**Fig. 1.**
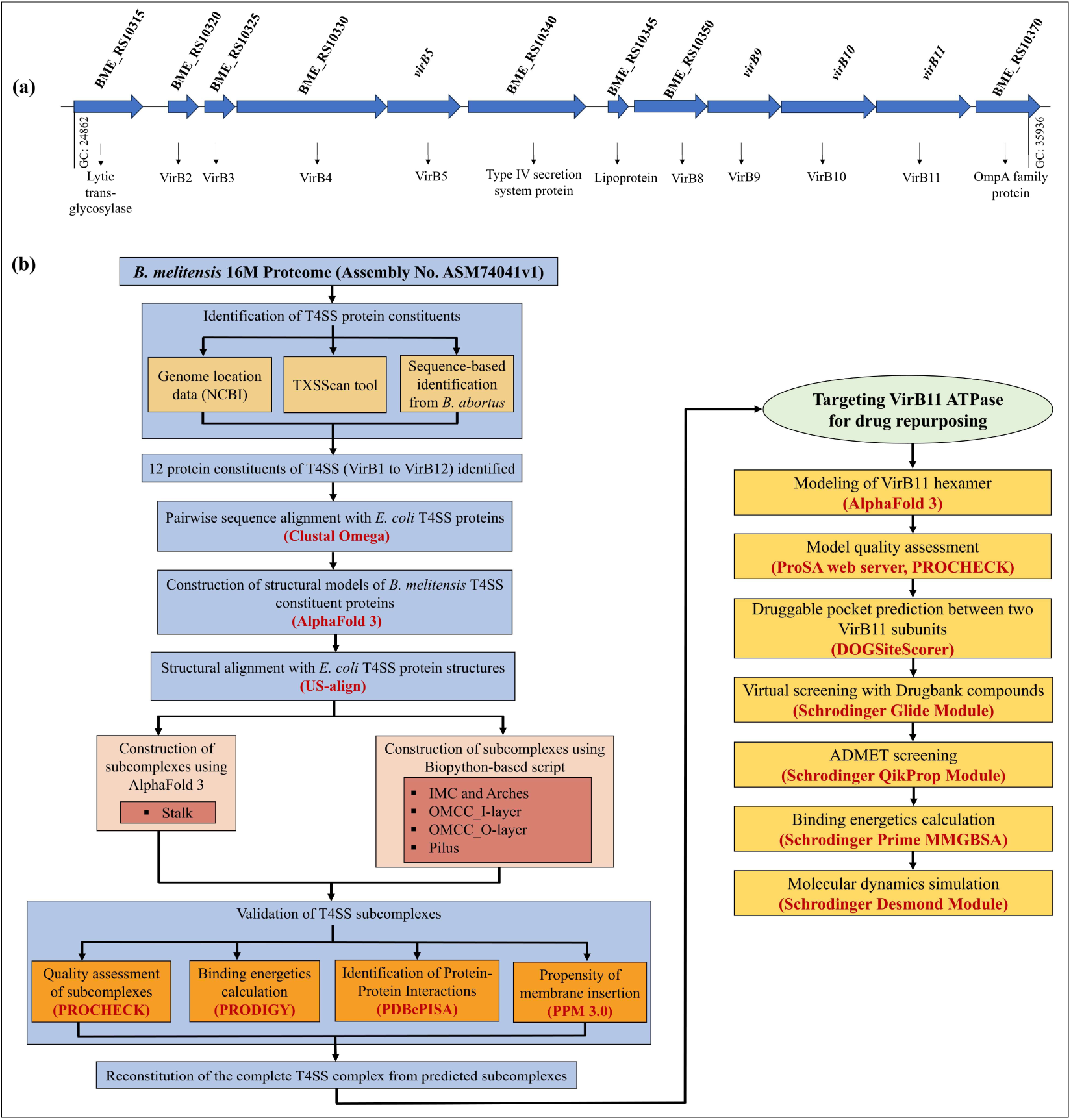
**(a)** Schematic representation of the genomic organization of *virB* operon in *B. melitensis* as annotated in the NCBI database (GC represents Genomic Coordinates). **(b)** Computational pipeline for assembly of T4SS components, followed by structure-based virtual screening of compounds from DrugBank targeting the VirB11 dimeric interface.

### 2.2 Pairwise sequence alignment with *E. coli* T4SS homologs

Following the identification of all T4SS constituent proteins in Bm, pairwise sequence alignment was performed against their respective Ec homologs using Clustal Omega [39] (https://www.ebi.ac.uk/jdispatcher/msa/clustalo, accessed on 5 September 2025). This was done to assess the degree of sequence similarity between the two bacterial species (Table 1).

**Table 1:** Identification and comparison of *B. melitensis* T4SS proteins with *E. coli* homologs.

| *B. melitensis* |  |  |  |  | *E. coli* |  | Sequence<br>similarity<br>(%) | Structural<br>similarity<br>(US-align<br>RMSD (Å)) |
| --- | --- | --- | --- | --- | --- | --- | --- | --- |
| T4SS<br>proteins | NCBI accession<br>no. | NCBI Annotation | Method of<br>identification | pTM score of<br>AlphaFold 3<br>models | *E. coli* homologs<br>(PDB Ids) | NCBI accession<br>no. |  |  |
| VirB1 | WP_004680997.1 | Lytic transglycosylase<br>domain-containing<br>protein | TXSScan | 0.77 | TrwN (No crystal<br>structure) | WP_032491146.1 | 24.2 | - |
|  | WP_004681227.1 | Lytic transglycosylase<br>domain-containing<br>protein | TXSScan | 0.73 | TrwN (No crystal<br>structure) | WP_032491146.1 | 44.15 | - |
| VirB2 | WP_002967165.1 | TrbC/VirB2 family<br>protein | Gene location,<br>TXSScan | 0.45 | TrwL (8S6H) | WP_031942907.1 | 20.83 | 2.93 |
| VirB3 | WP_002966512.1 | Type IV secretion<br>system protein VirB3 | Gene location,<br>TXSScan | 0.48 | TrwM (7O41) | WP_012443546.1 | 32.69 | 3.02 |
| VirB4 | WP_004681223.1 | VirB4 family type IV<br>secretion system<br>protein | Gene location,<br>TXSScan | 0.91 | TrwK (7O41) | WP_250693943.1 | 43.8 | 5.08 |
| VirB5 | WP_002966514.1 | P-type DNA transfer protein VirB5 | Gene location, TXSScan | 0.53 | TrwJ (8RT9) | WP_032490329.1 | 30.32 | 4.63 |
| VirB6 | WP_004686828.1 | Type IV secretion system protein | TXSScan | 0.68 | TrwI (8RT9) | WP_032490330.1 | 25.39 | 3.65 |
| VirB7 | WP_002966516.1 | Lipoprotein | Sequence similarity with *B. abortus* homolog | 0.26 | TrwH (8RT6, 8RT4) | WP_012196427.1 | 31.91 | 2.41 |
| VirB8 | WP_002966517.1 | VirB8 family protein | Gene location, TXSScan | 0.62 | TrwG (7O41) | WP_032490331.1 | 40.87 | 3.05 |
| VirB9 | WP_002966518.1 | P-type conjugative transfer protein VirB9 | Gene location, TXSScan | 0.69 | TrwF (8RT6, 8RT4, 8RT5) | WP_032490332.1 | 36.36 | 3.11 |
| VirB10 | WP_004685137.1 | Type IV secretion system protein VirB10 | Gene location, TXSScan | 0.53 | TrwE (8RT6, 8RT4, 8RT5) | WP_032490333.1 | 35.85 | 2.58 |
| VirB11 | WP_005972055.1 | P-type DNA transfer ATPase VirB11 | Gene location | 0.92 | TrwD (No crystal structure) | WP_012196431.1 | 40.41 | - |
| VirB12 | WP_004681215.1 | OmpA family protein | Sequence<br>similarity with<br>*B. abortus*<br>homolog | 0.59 | - | - | - | - |

– Corresponding structural or sequence homologs for these constituent proteins are not available in *E. coli*.

### 2.3 Generation of structural models of T4SS proteins

Three-dimensional models of identified proteins were generated from AlphaFold 3 (https://alphafoldserver.com/, accessed on 11 September 2025) [40]. AlphaFold 3 provided several confidence metrics, including the pLDDT (predicted Local Distance Difference Test), PAE (Predicted Aligned Error), pTM (predicted Template Modeling), and ipTM (inter-chain predicted TM) scores, which were used to assess the prediction reliability [41]. Structural model of each protein was visualized in PyMOL 3.0 using atomic coordinate file [42]. AlphaFold 3 predicted models of monomeric proteins were used for further study.

### 2.4 Structural comparison of *B. melitensis* T4SS monomers with *E. coli* T4SS crystal structures

Atomic coordinate files of Ec T4SS protein structures were downloaded from RCSB PDB database (accessed on 7 October 2025) with PDB Ids mentioned in Table 1. Structural similarity among the Bm and Ec T4SS constituent proteins was assessed using US-align [43] online web server (https://aideepmed.com/US-align/, accessed on 12 October 2025) and the corresponding RMSD (Root Mean Square Deviation) values were considered to validate the predicted architecture [44]. A low RMSD value reflects greater confidence in the accuracy and reliability of the predicted structures. As all the aligned structures had RMSD < 10 Å, we used the AlphaFold 3 models for construction of Bm T4SS (Table 1).

### 2.5 Construction of T4SS protein subcomplexes

The complete T4SS was divided into five subcomplexes based on their spatial position within the bacterial envelope from inside to outside: inner membrane complex (IMC) and arches; the stalk; outer membrane core complex I-layer (OMCC I-layer); outer membrane core complex O-layer (OMCC O-layer); and the pilus [20]. Initially, AlphaFold 3 was used that predicted the stalk subcomplex with reliable pTM and pLDDT scores. However, the same approach could not construct rest of the subcomplexes with reliable scores. Therefore, an alternate approach was used to assemble the monomeric protein models into subcomplexes using crystal structures of Ec subcomplexes as template. A Biopython-based [45] script was used to assemble all the T4SS constituent proteins into subcomplexes based on Ec T4SS architecture. The script aligned and assembled multiple protein subunits into a single subcomplex using a reference structure that served as the template for spatial arrangement and orientation. The modeled subcomplexes were visualized in UCSF ChimeraX 1.9 using atomic coordinate files [46].

### 2.6 Quality assessment, energetics calculations, and protein-protein interactions

Overall quality of each modeled subcomplex was assessed using PROCHECK [47] available on SAVES v6.0 server (https://saves.mbi.ucla.edu/). PROCHECK provides a detailed check on the stereochemistry of a protein structure based on Ramachandran plot. The structural model of subcomplexes were further validated by calculating interaction energetics using the PRODIGY (PROtein binDIng enerGY prediction) server [48] (https://rascar.science.uu.nl/prodigy/), which provides Gibbs free energy (ΔG, kcal mol^-1^) or dissociation constant (K_d_, M) for intermolecular interactions. PRODIGY predicts the binding affinity (ΔG) of protein-protein complexes based on intermolecular contacts at the interface, typically within a 5.5 Å distance cutoff. The formula correlates the number of interatomic contacts (ICs) at the interface with experimental binding affinity, classified by polar, apolar, or charged atom types. Additionally, amino acid residues involved in PPI, including hydrogen bonds, salt bridges, and hydrophobic interactions, were analyzed using PDBePISA [49] (https://www.ebi.ac.uk/pdbe/pisa/, all tools accessed on 15 October 2025). PISA (Protein Interfaces, Surfaces and Assemblies) estimates energetics by analyzing solvation free energy changes, buried surface area, hydrogen bonds, and salt bridges upon complex formation. All the parameters were also evaluated for crystal structure of Ec subcomplexes which served as positive control for validation [50].

### 2.7 Validation of membrane insertion of membrane embedded subcomplexes

The PPM (Positioning of Proteins in Membranes) 3.0 web server [51] was used to assess the potential of IMC and OMCC O-layer to insert into the inner and outer membrane, respectively (https://opm.phar.umich.edu/ppm_server3_cgopm, accessed on 28 October 2025). The PPM 3.0 webserver determines the number of amino acid residues embedded in membranes or micelles. It also evaluates protein stability within the membrane by calculating the ΔG transfer energy, which represents the energy required to move the protein from an aqueous environment into the lipid bilayer. The PPM 3.0 server inserts the protein into the inner membrane composed of phosphatidylethanolamine, phosphatidylglycerol, and cardiolipin (PE:PG:CL), while the outer membrane consists of PE:PG:CL in the inner leaflet and lipopolysaccharide (LPS) in the outer leaflet. The IMC and OMCC O-layer of Ec were also evaluated for membrane insertion as positive controls.

### 2.8 Homology modeling and model quality assessment of VirB11 ATPase

VirB11 is a hexameric ATPase present in cytoplasm of *Brucella* spp. [27]. Homology modeling of hexameric VirB11 of Bm was performed using AlphaFold 3 (accessed on 11 September 2025). Six copies of amino acid sequence of VirB11 were submitted to AlphaFold 3 server. Similarly, dimer of VirB11 was constructed using AlphaFold 3 for virtual screening of FDA-approved compounds against its dimeric interface. The monomeric structure of VirB11 was structurally aligned with monomeric crystal structure of *B. suis* VirB11 ATPase (PDB Id: 2GZA) using US-align online web server and structural similarity was assessed based on RMSD value. Furthermore, VirB11 homology was checked with the human proteome (TaxID: 9606) using BLASTP (E-value < 1.0e-03, bitscore > 100) (https://blast.ncbi.nlm.nih.gov/Blast.cgi?PROGRAM=blastp&PAGE_TYPE=BlastSearch&LINK_LOC=blasthome) (accessed on 08 December 2025). Following structural modeling, the quality of Bm VirB11 structural model was assessed using ProSA-web server (https://prosa.services.came.sbg.ac.at/prosa.php) [52] and PROCHECK (https://saves.mbi.ucla.edu/) [47] (both accessed on 9 December 2025). The ProSA (Protein Structure Analysis) web server compares the input protein model with the known experimentally determined structures available in the PDB and generates a Z-score. PROCHECK was used for Ramachandran plot analysis. Structural stability and stereochemical quality of the predicted model was validated based on Z-score and the distribution of amino acid residues in the Ramachandran plot.

### 2.9 Prediction of Druggable Pocket at VirB11 dimeric interface

DOGSiteScorer (https://proteins.plus/help/dogsite) (accessed on 9 December 2025) [53] was used to identify the druggable pocket in the VirB11 dimer. PDB coordinate file of dimeric VirB11 was used as input and potential binding pockets, along with their corresponding druggability scores, were predicted using a support vector machine (SVM)-based approach by DOGSiteScorer. The druggability score ranges from 0 to 1, where higher values indicate a greater likelihood that the pocket is druggable [53].

### 2.10 Virtual screening of compounds against predicted VirB11 druggable pocket

FDA approved drugs available at DrugBank database (https://go.drugbank.com/) were used for virtual screening (accessed on 10 December 2025) [54]. Structural data (.sdf) file of three-dimensional structures of 2648 drug compounds was downloaded from DrugBank. Virtual screening was performed using Schrodinger suite 2025-3 [55]. Initially, the structural model of VirB11 dimer was prepared using the Protein Preparation Wizard in Schrodinger suite 2025-3, followed by energy minimization using the OPLS4 force field. Interacting amino acid residues listed in Figure 7 were specified to generate receptor grid around predicted druggable pocket. Ligand molecules were prepared for docking using the LigPrep module of Schrodinger suite 2025-3, with structures provided in .sdf format. For virtual screening, the prepared ligand molecules were docked onto the receptor grid of prepared VirB11 dimer using Glide module [56] of the Schrodinger suite 2025-3. It employs an algorithm to identify favorable hydrogen-bonding, hydrophobic, and electrostatic interactions while penalizing steric hindrance. All docked poses are re-evaluated and ranked after energy minimization using GlideScore scoring function [56]. Ligand molecules with docking score ≤ - 7.0 kcal/mol were selected for ADMET (adsorption, distribution, metabolism, excretion, and toxicity) screening.

### 2.11 ADMET Screening

The ADMET properties of the selected ligands, having docking scores ≤ -7.0 kcal/mol, were evaluated using the QikProp module of the Schrodinger suite 2025-3, followed by their filtration. Nine criteria were applied for screening: molecular weight < 500 Da, solvent accessible surface area (SASA, 300-1000 Å), one violation of Lipinski’s Rule of Five, ≥80% predicted human oral absorption, QPPCaco ≥ 500, QPlogHERG < –5, donor HB ≤ 5, acceptor HB ≤ 10, and QPlogPo/w (-2 to 6.5). The QikProp user manual was used to determine acceptable ranges for each property [57]. Finally, three protein-ligand complexes were filtered for further assessment of binding energetics and molecular dynamics simulation.

### 2.12 Calculation of Binding Free Energy

Prime module of the Schrodinger suite 2025-3 was used to calculate the binding free energies (ΔG_bind_) of the selected protein-ligand complexes through the molecular mechanics– generalized born surface area (MM–GBSA) approach. The method employs the optimized potentials for liquid simulation (OPLS4) force field and variable solvent generalized born (VSGB) solvent model, along with search algorithms [58]. The binding free energy was determined using the following relationship:

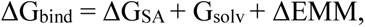

where ΔG_SA_ depicts the change in surface area energies between the protein-ligand complexes and the individual protein and ligands, ΔG_solv_ represents the difference in the solvation energies of the complexes and the individual protein and ligands, and ΔEMM reflects the difference in molecular mechanics energies between the protein-ligand complexes and the individual protein and ligands when minimized.

### 2.13 Molecular Dynamics Simulation

The structural stability of the selected protein-ligand complexes was assessed by performing molecular dynamics simulations (MDS) using the Desmond module in the Schrodinger suite 2025-3. The docked protein-ligand complexes were first prepared using the System Builder, where each protein-ligand complex was solvated with TIP3P water molecules and enclosed with an orthorhombic simulation box. The OPLS4 force field was applied at physiological pH 7.4, and Na+ and Cl-ions were added to neutralize the system and maintain a salt concentration of 0.15M. After generation of a complete system, MDS was performed for 200 ns under NPT conditions at a bar pressure of 1.01325 and a constant temperature of 300K. The Simulation Interactions Diagram (SID) tool was subsequently used to analyze the trajectories, providing RMSD and root mean square fluctuation (RMSF) plots, and protein-ligand interaction profiles for each trajectory frame of the simulation.

## 3. Results

### 3.1 Identification of VirB proteins in *B. melitensis* proteome

The complete proteome of *B. melitensis* biovar 1 strain 16M, downloaded from NCBI, contains 2978 protein sequences in FASTA format. The Bm T4SS is composed of 12 VirB proteins as mentioned in literature but not all proteins are annotated as VirB family proteins (Figure 1a). Therefore, to identify the accession numbers of Bm T4SS constituent proteins from the complete proteome, three complementary approaches were employed. In the first approach, proteins annotated as VirB were identified based on genomic location data of the Bm T4SS operon available in NCBI. Eight proteins (VirB2, VirB3, VirB4, VirB5, VirB8, VirB9, VirB10, and VirB11) were found to be annotated as VirB family proteins. Remaining unannotated proteins (VirB1, VirB6, VirB7, and VirB12) were inferred based on their neighbouring annotated proteins. The region upstream of *virB2* was likely to encode VirB1 (WP_004681227.1), and the genes located between *virB5* and *virB8* were likely to encode VirB6 (WP_004686828.1) and VirB7 (WP_002966516.1). Similarly, the gene downstream of *virB11* was likely to encode VirB12 (WP_004681215.1) (Table 1). To confirm the identification of these four unannotated T4SS constituent proteins, alternate approaches were used. The second approach employed TXSScan tool that leverages Hidden Markov Models (HMMs) to identify genes encoding proteins belonging to various bacterial secretion systems. The tool provides the accession no. of proteins, gene names, and corresponding E-values from the HMM or sequence alignment, indicating the significance of the match (E-value < 0.001) (Table S1). Complete proteome file of Bm was used as input file and TXSScan identified nine proteins (two homologs of VirB1, and one homolog of each VirB2, VirB3, VirB4, VirB5, VirB6, VirB8, VirB9, and VirB10) as VirB family proteins which are part of Bm T4SS. The accession number of the VirB1 (WP_004681227.1), and VirB6 (WP_004686828.1) proteins were consistent with those predicted from genome location-based annotation. In the third approach, the identity of VirB7, and VirB12 proteins was validated based on sequence similarity with *B. abortus* homologs. BLASTP analysis showed that the proteins WP_004681227.1 and WP_004681215.1 from Bm are homologous to the *B. abortus* VirB1 (AAF73894.1) and VirB12 proteins (AAF73905.1), with E-values of 2e-178 and 3e-125, respectively.

### 3.2 Assessment of sequence and structural homology with *E. coli* T4SS proteins

Pairwise sequence alignment of T4SS constituent proteins was carried out between the identified Bm proteins and their Ec homologs (Table 1). Except VirB1 (WP_004680997.1), VirB2 (WP_002967165.1), and VirB6 (WP_004686828.1), all other proteins exhibited >30% sequence similarity. Subsequently, structural models of Bm T4SS proteins available in the AlphaFold database were retrieved and remaining protein models were generated using AlphaFold 3. The resulting models were evaluated based on pTM and pLDDT scores (Table 1). A pTM score greater than 0.5 indicates an overall reliable structure. The predicted models met this threshold except VirB2, VirB3, and VirB7. Since pTM scores are less informative for small or short proteins, the pLDDT values are considered for such models. Notably, for VirB2, VirB3, and VirB7, the pLDDT scores were above 70 across most of the regions, indicating a reliable model. Structural alignment of these monomeric models with crystal structure of their corresponding *E. coli* homologs using US-align indicated a high overall structural similarity with low RMSD values ranging from 2.41 Å to 5.08 Å (Table 1).

### 3.3 Assembly of T4SS subcomplexes

VirB proteins were assembled into five T4SS subcomplexes: IMC and arches; the stalk; OMCC I-layer; OMCC O-layer; and the pilus. Initially, AlphaFold 3 was employed to predict the structures of T4SS subcomplexes. The stalk subcomplex was predicted with a reliable interface confidence, having an ipTM score of 0.7. However, the remaining subcomplex models exhibited ipTM scores below 0.6, indicating a low confidence in their inter-chain interactions. The inability to model large protein complexes is a major limitation of AlphaFold 3 [59]. To overcome this limitation, a Biopython-based script [45,60] was employed to assemble the remaining subcomplexes where the crystal structures of corresponding Ec T4SS subcomplexes were used as template.

Based on above predictions, we determined the structural organization of five T4SS subcomplexes in Bm. The innermost subcomplex, *IMC and arches*, is composed of VirB3, VirB4, and VirB8 (Figure 2). It contains six protomers arranged symmetrically around a central axis. A protomer represents a repeating multi-protein unit composed of multiple monomers assembled together. In IMC, each protomer includes one VirB3, two VirB4, and two VirB8 N-terminal tails. The overall architecture can be visualized as a six-spoke wheel, with each protomer acting as one spoke. VirB4 ATPase forms a double ring configuration (inner—outer) that is essential for powering substrate transfer and pilus assembly. VirB3 is a small membrane-associated protein that contributes to IMC stability, acting like a membrane anchor. N-terminal tails of VirB8, embedded within the inner membrane, are positioned at the interface between the IMC and periplasmic arches. The arches form a periplasmic ring-like assembly composed of periplasmic domains of VirB8. The arches sit directly above the IMC and connect it to the stalk and the OMCC subcomplex. Protruding from the IMC into the periplasm is *the stalk* subcomplex. It consists of pentamers of VirB5 and VirB6 which together form a central, cone-shaped structure that connects the IMC to the OMCC (Figure 3a). VirB5 and VirB6 form stacked, concentric pentameric rings that stabilize the central axis of T4SS. VirB6 forms the base that recruits VirB2 pilin subunits during pilus biogenesis, while VirB5 forms the upper cap of pilus that interacts with the host cell [25]. Connecting the stalk to the outer membrane is the *OMCC* (I-layer and O-layer) subcomplex. *OMCC I-layer* is formed by the N-terminal domains of VirB9 (VirB9_NTD_) and VirB10 (VirB10_NTD_) (Figure 4). The I-layer assembles into a 16-membered ring formed by repeating heterodimers (VirB9_NTD_-VirB10_NTD_). The I-layer provides a periplasmic scaffold that stabilizes the overall OMCC structure. (iv) *OMCC O-layer*: The O-layer is composed of 14 repeating subunits of heterotrimer-VirB7, C-terminal domains of each VirB9 (VirB9_CTD_), and VirB10 (VirB10_CTD_) (Figure 5). These subunits form a barrel-shaped outer membrane pore. VirB10_CTD_ spans the outer membrane to anchor the O-layer and VirB7 stabilizes the structure by interacting with both VirB9_CTD_ and VirB10_CTD_. The OMCC displays a pronounced asymmetry between its two layers: the 16-fold I-layer and the overlying 14-fold O-layer. In Ec, this asymmetry is thought to provide structural flexibility, allowing the complex to accommodate dynamic motions during pilus extrusion and substrate secretion [61]. Lastly, *the pilus* subcomplex is primarily composed of VirB2 subunits (Figure 3b). VirB2 is major pilin protein that polymerizes into a columnar pilus filament which is assembled as a five-start helical filament at VirB5-VirB6 interface. Studies in Ec have shown that during pilus biogenesis, five VirB2 subunits first assemble into a pentameric ring that matches the five-fold symmetry of VirB6 stalk. As new VirB2 layers polymerize beneath the previous one, the growing pilus pushes the VirB5 pentamer upward and outward forming the distal cap of the pilus [25].

**Fig. 2.**
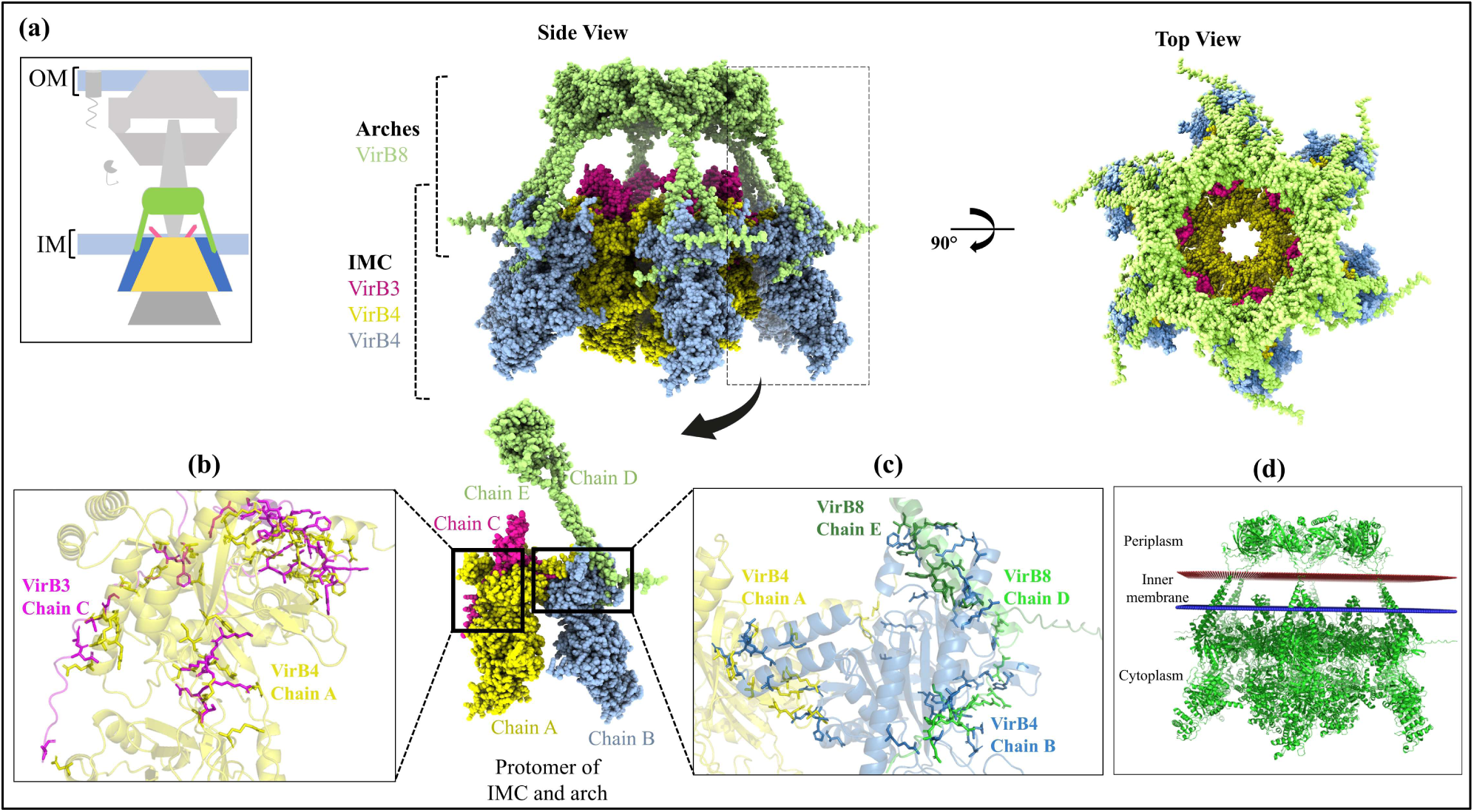
Structural organization of the Inner Membrane Complex (IMC) and Arches of the T4SS in *B. melitensis*. **(a)** Side and top views of the modeled IMC and arches subcomplex are shown highlighting the ring-like arrangement. VirB4 ATPase has double-layered hexameric rings: inner and outer ring. The tails from hexamer of dimeric VirB8 is connected to the outer ring while the periplasmic domains of VirB8 constitutes the base of the periplasmic arch. Enlarged view of a single protomer comprising VirB4_inner_ (chain A, yellow), VirB4_outer_ (chain B, blue), VirB3 (chain C, magenta), and VirB8 (chain D, E, light green) is shown. **(b)** The key amino acid residues involved in hydrogen bonding between VirB3-VirB4 are depicted. **(c)** The amino acid residues involved in hydrogen bonding between VirB4_inner_ – VirB4_outer_, and VirB4_outer_ – VirB8 dimeric tails are highlighted. **(d)** Propensity of membrane insertion of Bm IMC and Arches subcomplex as predicted by PPM 3.0 server.

**Fig. 3.**
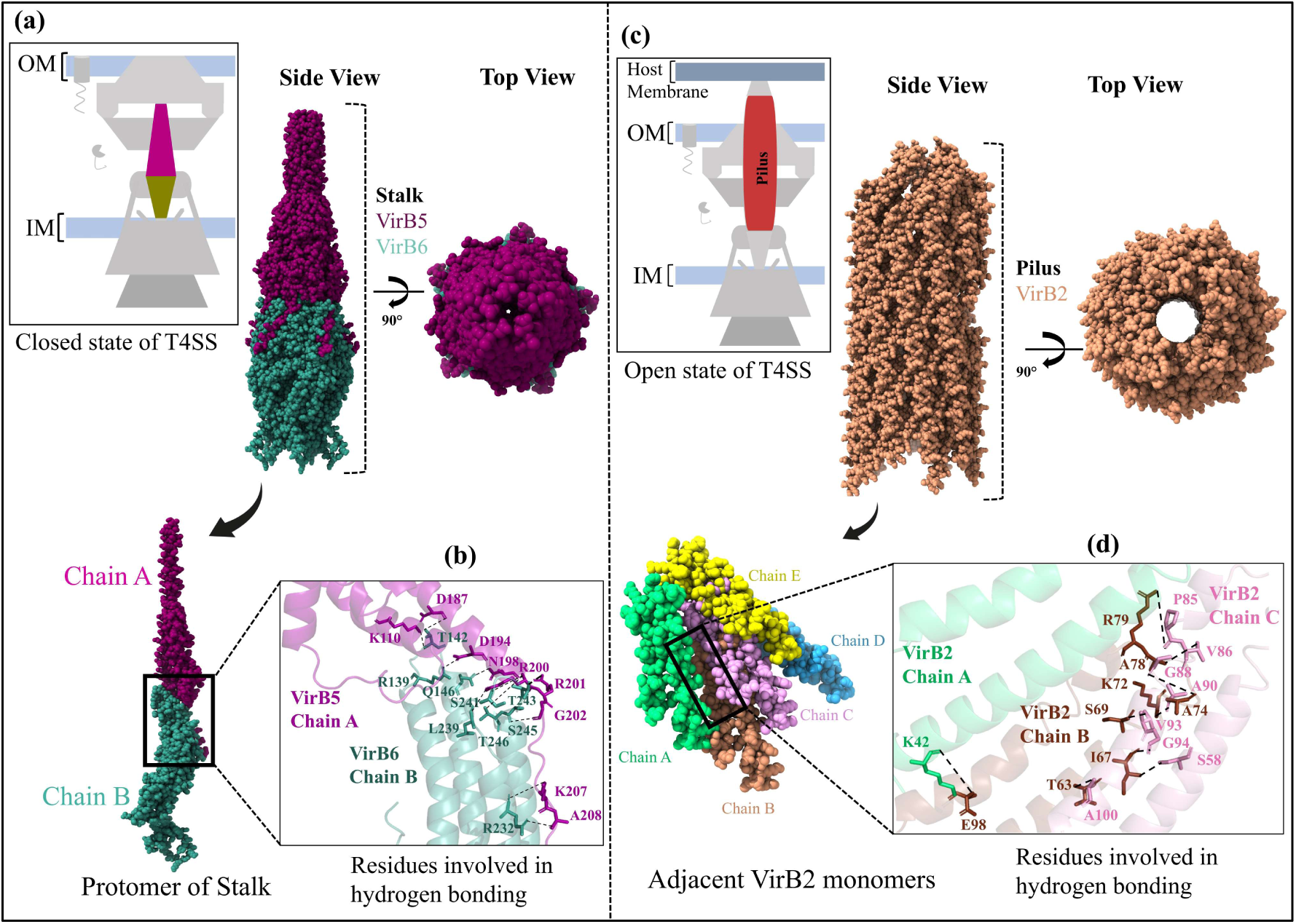
Structural arrangement of the stalk and the pilus component T4SS in *B. melitensis*. The stalk and pilus form a continuous conduit that spans the periplasm connecting the IMC to the OMCC. **(a)** The stalk complex comprises VirB5 (chain A, dark purple), and VirB6 (chain B, cyan) subunits that assemble into a pentameric stalk. **(b)** The panel shows the key hydrogen bond interactions between VirB5-VirB6. **(c)** The pilus is composed of VirB2 subunits, forming a spiral filament extending from the outer membrane surface. The side and top views reveal a hollow, cylindrical arrangement consistent with secretion pili. **(d)** Each VirB2 subunit interacts with adjacent monomers (chains A–E), stabilized by hydrogen bonds displayed in the panel.

**Fig. 4.**
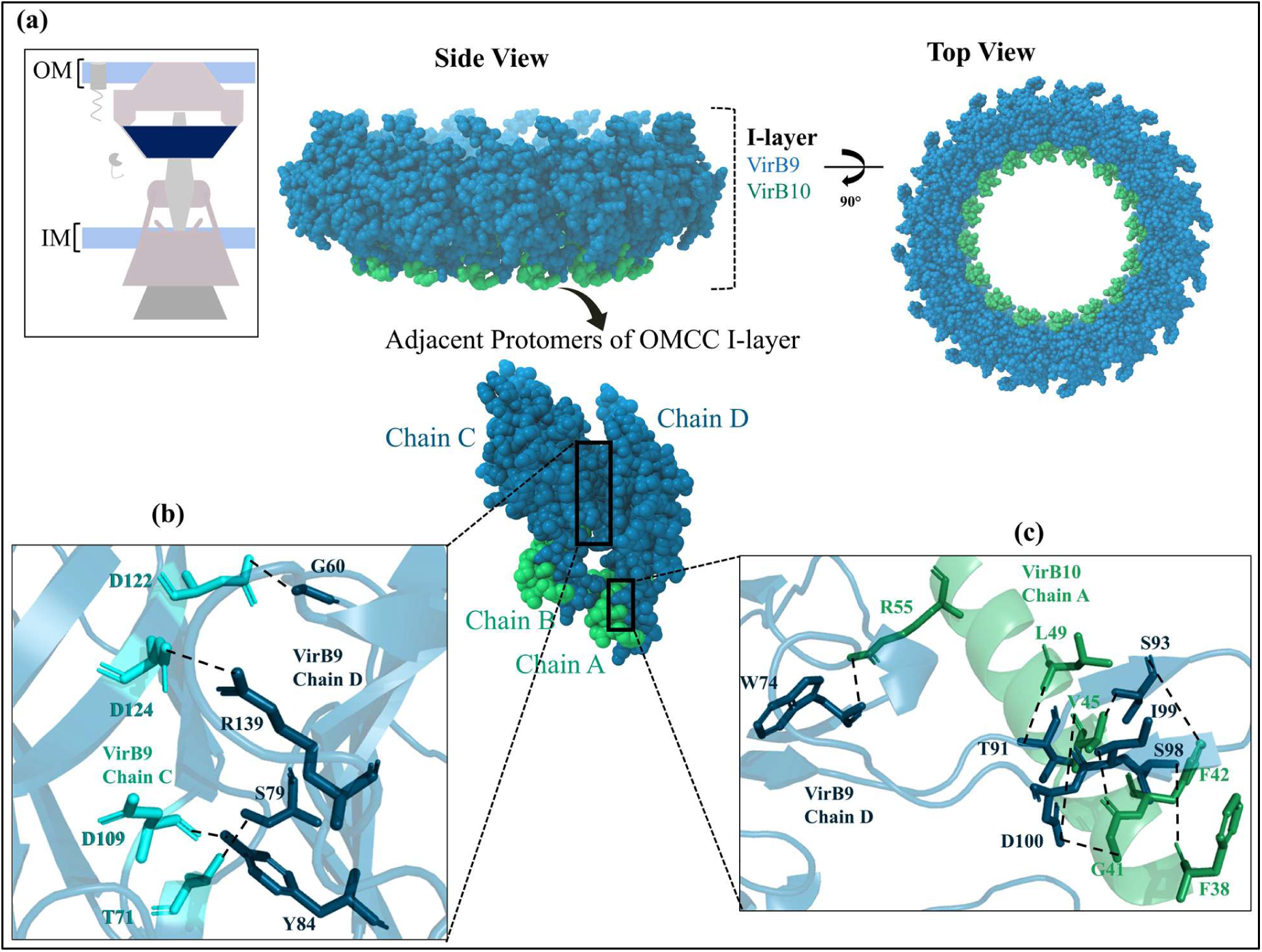
Structural organization of the Outer Membrane Core Complex (OMCC) I-layer of the T4SS in *B. melitensis*. The predicted OMCC I-layer subcomplex exhibits a barrel-shaped architecture located beneath the O-layer. **(a)** The side view displays the dense, barrel-like arrangement of the complex, while the top view reveals a 16-fold symmetrical assembly surrounding a central channel. The structure is composed of VirB9 (chain C, D, blue), and VirB10 (chain A, B, green), that oligomerize into the complete I-layer ring. **(b)** Hydrogen bond interactions stabilizing protein-protein interfaces between two adjacent VirB9_NTD_ are shown. **(c)** Key amino acid residues forming hydrogen bonds between VirB9_NTD_-VirB10_NTD_ are highlighted.

**Fig. 5.**
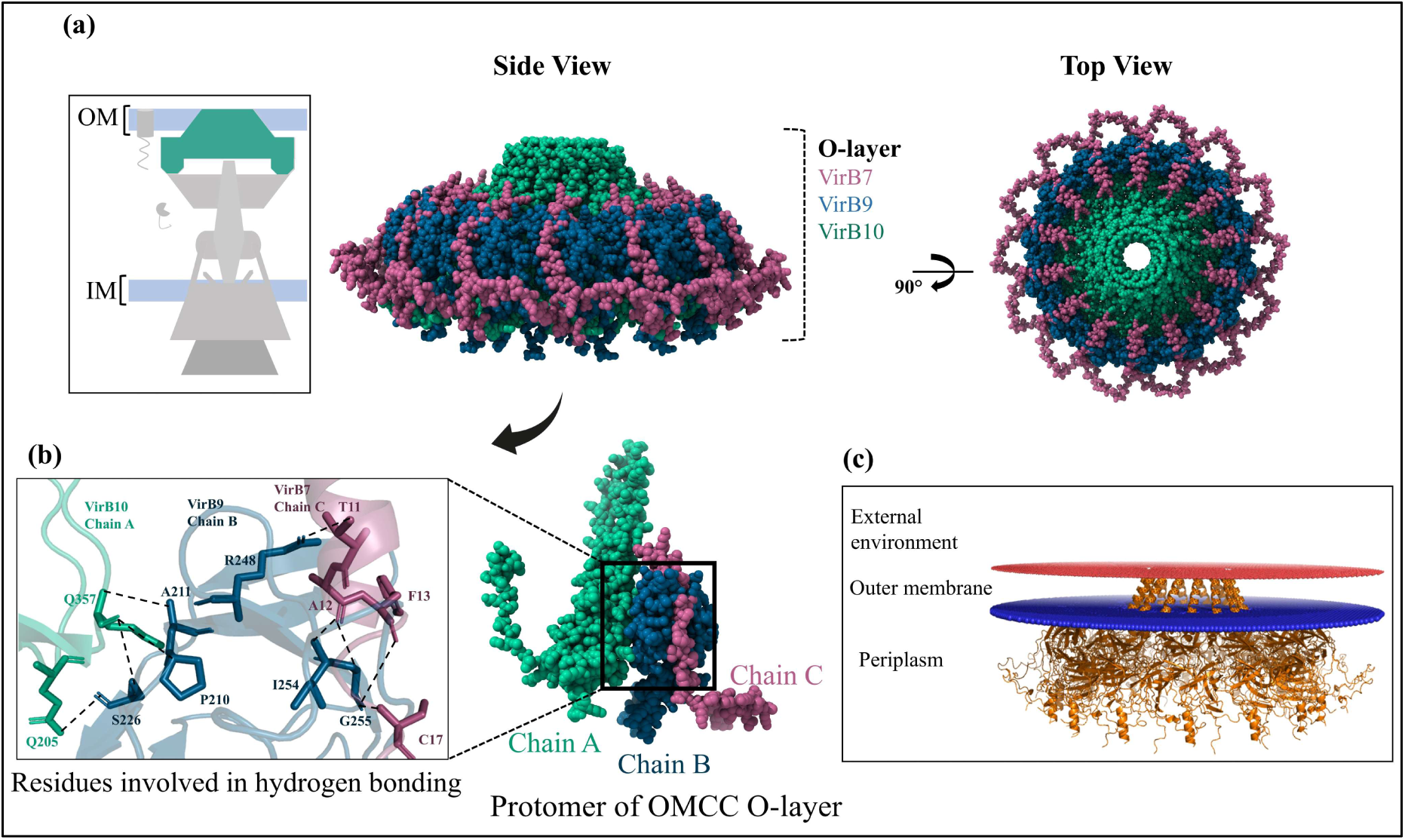
Structural architecture of the Outer Membrane Core Complex (OMCC) O-layer of Bm T4SS. The modeled OMCC_O-layer shows a 14-fold arrangement comprising VirB9 (chain B, blue), VirB10 (chain A, green), and VirB7 (chain C, purple). All three monomers form a stable trimeric protomer unit that oligomerizes to produce the full O-layer ring. **(b)** The panel highlights the key amino acid residues involved in hydrogen bonding between the subunits VirB7-VirB9-VirB10. **(c)** Membrane insertion propensity of OMCC O-layer predicted by PPM 3.0 server.

Three VirB proteins — VirB1, VirB11, and VirB12 — perform important functional roles despite not being incorporated into any T4SS subcomplex. VirB1 is a periplasmic lytic transglycosylase enzyme that remodels the peptidoglycan layer to facilitate the assembly of T4SS subcomplexes [23]. VirB11 is a cytoplasmic hexameric ATPase that interacts transiently with the IMC VirB4 ATPase to regulate energy-dependent steps [27]. VirB12 is a surface-associated accessory protein that is strongly expressed during infection and serves as an immunogenic biomarker for disease diagnosis [62].

The above subcomplexes and three VirB proteins (VirB1, VirB11, and VirB12) were then assembled manually to predict the overall architecture of Bm T4SS (Figure 6).

**Fig. 6.**
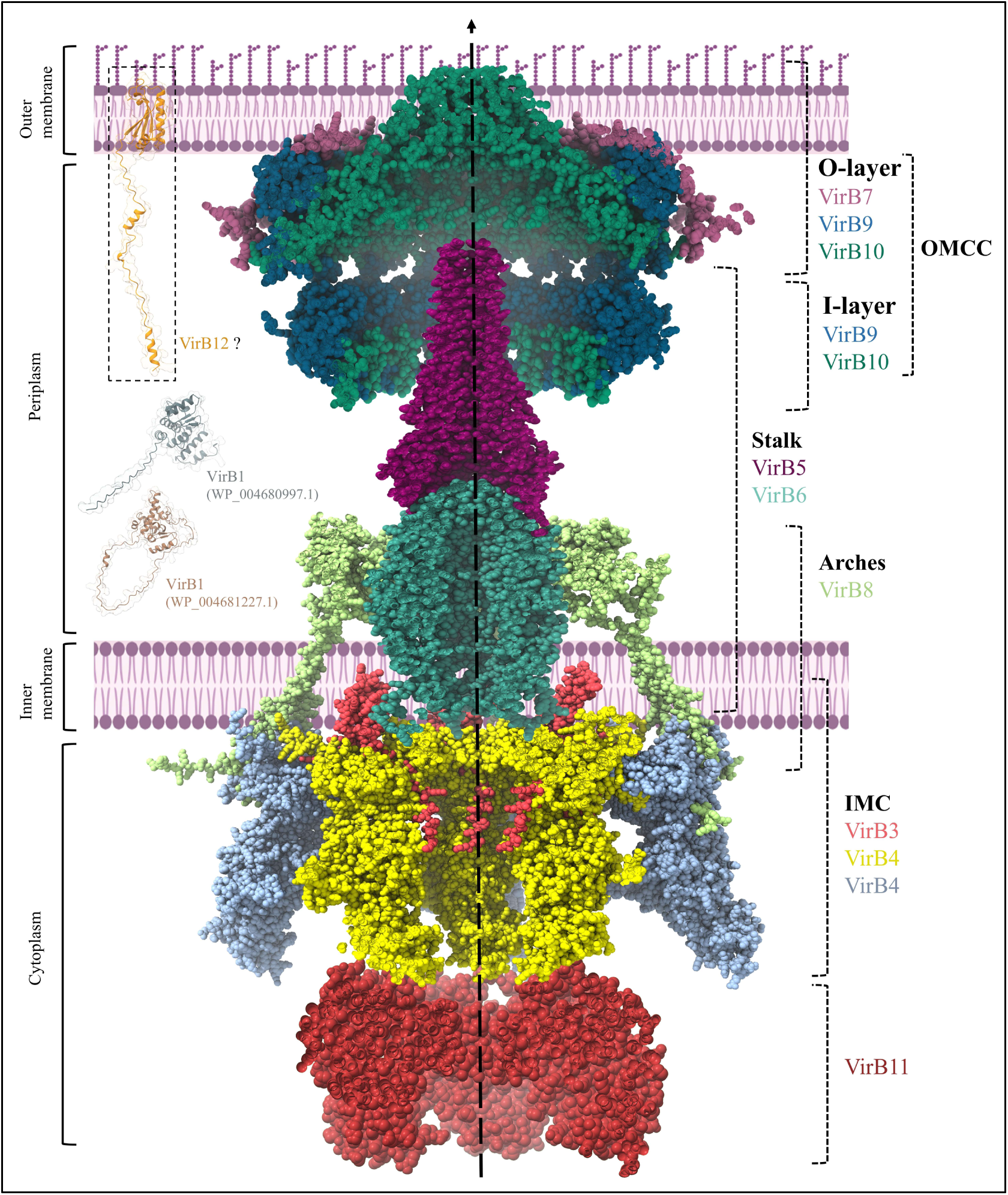
Complete architecture of the Type IV Secretion System in *B. melitensis*. The Bm T4SS is organized into five major subcomplexes-IMC and Arches, stalk, OMCC I-layer, OMCC O-layer, and pilus. The figure shows a closed state of T4SS without pilus. The IMC comprises VirB3, VirB4, and VirB8 subunits where VirB11 is coupled to VirB4 ATPase. The arches are formed by the periplasmic domains of VirB8. The stalk region is composed of VirB5 and VirB6, forming a continuous conduit. The OMCC presents a two-layered architecture: the O-layer (VirB7, VirB9_CTD_, and VirB10_CTD_), and the I-layer (VirB9_NTD_ and VirB10_NTD_). Other components include two predicted lytic transglycosylases (VirB1) within the periplasm; and surface-exposed VirB12.

### 3.4 Quality assessment and binding energetics calculation

Further, the predicted models of Bm T4SS subcomplexes were validated using crystal structure of Ec T4SS subcomplexes as positive control. The Bm T4SS subcomplexes were evaluated using PROCHECK to assess the stereochemical quality based on Ramachandran plot statistics. Typically, a good-quality model should have ≥90% of the amino acid residues in the favoured region. In the Ramachandran plot analysis, IMC and arches had 93.9% residues, the stalk had 93.2% residues, OMCC I-layer had 93.8% residues, OMCC O-layer had 87.6%, and the pilus had 98.3% residues falling within the favoured regions (Figure S1). Thus, all subcomplex models exceed the threshold, except for the OMCC O-layer, indicating that the predicted structures are stereo-chemically reliable.

PRODIGY server predicted the ΔG (kcal mol⁻¹) and K_d_ values (M) between adjacent monomers of each T4SS subcomplex. A more negative ΔG value indicates a stronger and more stable binding interaction. Also, a lower K_d_ value (e.g., nanomolar or picomolar range) indicates a stronger binding affinity, while a higher value (e.g., micromolar or higher) indicates a weaker interaction. Since crystal structures of Ec T4SS subcomplexes are well established, they were used as positive control to define a meaningful comparison threshold. Four subcomplexes of Bm T4SS — the stalk, OMCC I-layer, OMCC O-layer, and the pilus — exhibited binding energetics similar to those of Ec T4SS (Table 2). The subcomplex of IMC and arches demonstrated stronger interaction parameters than their Ec counterparts. The ΔG (-51.5 kcal mol^-1^) and K_d_ values (1.6×10^−38^ M) for the VirB3-VirB4 interaction in Bm IMC were notably more negative than those observed in Ec IMC (ΔG = -20.3 kcal mol^-1^, K_d_ = 1.2×10^−15^ M). These values signify stronger and thermodynamically stable interactions within the IMC complex.

**Table 2:**
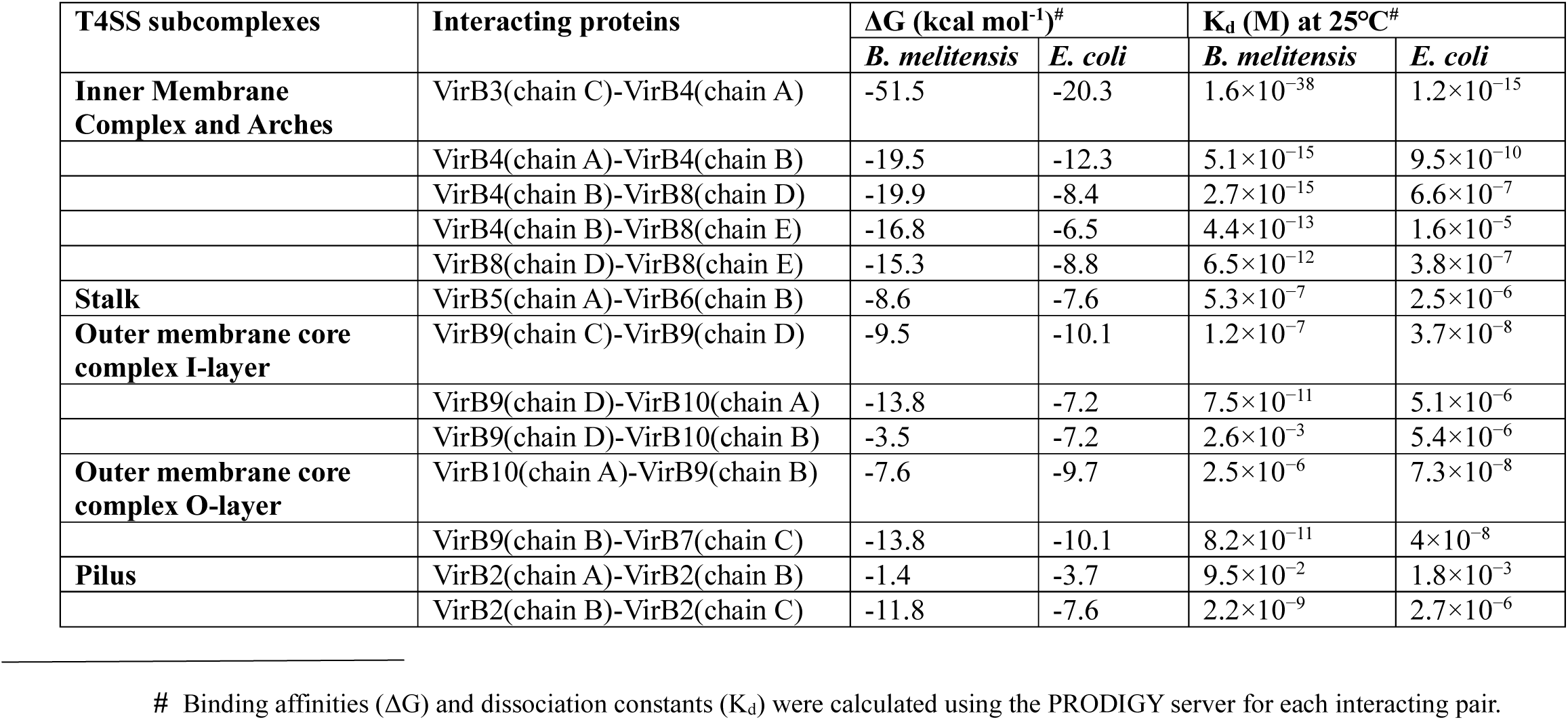
Binding affinity analysis of T4SS subcomplex interactions in *E. coli* and *B. melitensis*.

### 3.5 Analysis of Protein–Protein Interaction interfaces

Intermolecular interactions between adjacent monomers of subcomplexes were assessed to confirm the thermodynamically stable assemblies. PDBePISA revealed the hydrogen bonds, salt bridges, and hydrophobic contacts among the interfacing residues along with ΔG values. All interacting protein pairs exhibited negative ΔG values, indicating energetically favourable and stable interactions, except for the adjacent VirB4 monomers located in the inner and outer rings of the IMC, which showed a slightly positive ΔG (0.4 kcal mol^-1^). ΔG represents the sum of both stabilizing (hydrogen bonds, salt bridges, van der Waals, and hydrophobic surfaces) and destabilizing (desolvation energy, electrostatic repulsion, entropic costs) forces, so a positive value can still arise. However, the presence of 34 residues forming hydrogen bonds and salt bridges indicates significant interface interactions. The key amino acid residues participating in hydrogen bond formation were identified and mapped within each subcomplex, as illustrated in Figures 2, 3, 4, and 5; Table 3.

**Table 3:** Interfacial residue analysis of T4SS subcomplexes in *B. melitensis* predicted by PDBePISA.

| T4SS subcomplexes | Interacting proteins | $\Delta G$ (kcal mol<sup>-1</sup>)<sup>#</sup> | | Interfacing residues<sup>†</sup> involved in hydrogen bonding in *B. melitensis* | Interfacing residues<sup>&</sup> involved in salt bridge formation in *B. melitensis* |
| --- | --- | --- | --- | --- | --- |
|  |  | *B. melitensis* | *E. coli* |  |  |
| **Inner Membrane Complex and Arches** | VirB3(chain C)-VirB4(chain A) | -16.9 | -27.4 | **Chain C-** T2, N9, A10, S12, A13, G14, F21, C24, R79, K84, Q86, F87, R88, R88, I90, H91, F92, N93, R94, R100, A101, S102, A103, R112, K113, R114, N9, S12, G14, I20, F21, G82, Q86, H91, T95, W99, S102, Y104, K111, E115, S116 | **Chain C-** M1 |
|  |  |  |  | **Chain A-** D740, E56, L59, K62, D63, Y257, D253, Y385, N382, Y385, E14, P19, D223, R15, D223, C220, Q10, E14, V231, Y376, N405, N120, S119, T88, R61, Q64, N396, A390, R388, R384, Y385, F20, W230, T228, Y232, Y376, L17, A18, Y123, I45, N405, A408, D179, R411 | **Chain A-** D740 |
|  | VirB4(chain A)-VirB4(chain B) | 0.4 | -13.2 | **Chain A-** Y7, V215, R216, A195, S219, G28, S25, P29, T188, T31, H193, S5, E190, D191, C197 | **Chain A-** R216 |
|  |  |  |  | **Chain B-** A233, D256, A258, N259, Q297, L301, Q302, A303, T304, E305, D306, N306, N360, R15, N259, H260, Q311 | **Chain B-** D256 |
|  | VirB4(chain B)-VirB8(chain D) | -3.4 | × | **Chain B-** R61, R174, I183, I45, R44, R181, S187, V217, C220, T170, E169, D179, Y123, E202, Y189, A195, E224 | **Chain B-** E121, E224 |
|  |  |  |  | **Chain D-** Q6, S7, Q9, S11, V12, N14, D24, E25, P8, K10, S11, K13, Q16, N18, A19, A32 | **Chain D-** K13, R35 |
|  | VirB4(chain B)-VirB8(chain E) | -13.9 | × | **Chain B-** R159, R145, V143, N140, V141, Q154, L156, E153, E149, D36, F139 | **Chain B-** E149 |
|  |  |  |  | **Chain E-** P20, W29, A32, H33, S21, Y23, E25, N28, R35, V34, V37 | **Chain E-** R35 |
|  | VirB8(chain D)-VirB8(chain E) | -30.9 | × | **Chain D-** I62, G59, A63, M65, K69, G174, V101, S97, D103, Q109, L58, G64 | - |
|  |  |  |  | **Chain E-** I54, T55, L57, L58, A63, E93, S95, V96, Y98, I60, G61, I62, S97 | - |
| **Stalk** | VirB5(chain A)-VirB6(chain B) | -4.2 | -8.4 | **Chain A-** K110, N198, R200, R201, D187, D194, G202, K207, A208 | **Chain A-** D194 |
|  |  |  |  | **Chain B-** T142, Q146, L239, S241, T243, T246, R139, S245, R232 | **Chain B-** R139 |
|  | VirB5(chain A)-VirB5(chain C) | -28.6 | -50.1 | **Chain A-** D1, G15, Q22, Q29, Q36, V41, Y92, K118, Q155, N162, N8, E14, G146, N143, E136, S122, S124, D126, T145, T182, D169, Y125, E183, Q176 | **Chain A-** D1, K118, E14, E136, D126, E136, D129, D169, E183 |
|  |  |  |  | **Chain C-** D1, Q15, Q22, Q29, L26, Q33, E32, E192, E68, K118, E152, Q159, H125, G42, N43, G47, E56, R64, Y149, N167, R171, H173, L177, K184 | **Chain C-** D1, E68, H125, R51, R56, R64, R171, H173, K184 |
|  | VirB6(chain B)-VirB6(chain D) | -69.3 | -30.4 | **Chain B-** N28, Y21, S212, R204, Y44, G286, K70, S67, Q63, M32, M24, D137, Q215, P197, P334, E58, N345 | - |
|  |  |  |  | **Chain D-** A102, F103, Y270, I273, S276, T292, K294, S295, G297, T301, F99, N106, S149, S279, S302, A335, Q343, R346 | - |
| **Outer membrane core complex_I-layer** | VirB9(chain C)-VirB9(chain D) | -1.8 | -7.2 | **Chain C-** G60, S79, Y84, R139 | **Chain C-** R54, R139 |
|  |  |  |  | **Chain D-** D122, T71, D109, D124 | **Chain D-** D109, D124 |
|  | VirB9(chain D)-VirB10(chain A) | -6.3 | -2.8 | **Chain D-** S98, I99, D100, S93, T91, W74 | - |
|  |  |  |  | **Chain A-** F38, G41, F42, V45, L49, R55 | - |
|  | VirB9(chain D)-VirB10(chain B) | -1.4 | -4.8 | - | - |
|  |  |  |  | - | - |
| **Outer membrane core complex_O-layer** | VirB10(chain A)-VirB9(chain B) | -3.9 | -0.1 | **Chain A-** Q357, Q205 | - |
|  |  |  |  | **Chain B-** A211, S226, P210 | - |
|  | VirB9(chain B)-VirB7(chain C) | -14.9 | -4.1 | **Chain B-** G255, R248, I254 | - |
|  |  |  |  | **Chain C-** C17, T11, A12, F13 | - |
| **Pilus** | VirB9(chain B)-VirB7(chain C) | -14.9 | -4.1 | **Chain B-** G255, R248, I254 | - |
|  |  |  |  | **Chain C-** C17, T11, A12, F13 | - |
|  | VirB2(chain B)-VirB2(chain C) | -24.8 | -27.7 | **Chain B-** T63, S69, K72, A74, A78, R79, I67 | - |
|  |  |  |  | **Chain C-** A100, V93, A90, G88, V86, P85, S58, G94 | - |

### 3.6 Validation of membrane insertion of membrane embedded subcomplexes

To determine which portions of the Bm IMC and OMCC O-layer are embedded within the inner and outer membranes, the PPM server was employed. The Ec T4SS served as a positive control for the comparison. The PPM analysis provides the ΔG_transfer_ energy, which reflects the thermodynamic stability of membrane insertion. In Bm, the ΔG_transfer_ values of the IMC and OMCC O-layer were -177.8 kcal/mol and -80.7 kcal/mol, respectively, while for Ec, the corresponding values were -184.3 kcal/mol and -95.8 kcal/mol. These closely comparable values indicate that the membrane insertion and stability of the Bm subcomplexes are energetically similar to those of Ec, suggesting correctness of our predicted models. Moreover, the orientation of the subcomplexes within the membrane, as predicted by PPM server, was also found to be highly consistent between the two species, supporting the structural conservation and functional resemblance of their T4SS assemblies (Figure S2).

### 3.7 Homology Modeling and Model Quality Assessment of VirB11 ATPase

The structural model of the VirB11 hexamer, generated by AlphaFold 3, showed high confidence with pTM and ipTM score of 0.88 and 0.86, respectively (Figure 7). The structure of VirB11 consists of an N-terminal domain (NTD) and a C-terminal domain (CTD) connected with a short linker region [27]. The monomeric model of Bm VirB11 was superimposed onto the crystal structure of *B. suis* VirB11 (PDB ID: 2GZA), yielding an RMSD value of 0.84 Å, which supports the reliability of the predicted model. The quality of the model was evaluated using the ProSA-web server where a Z-score of -10.47 indicated concordance with those of experimentally determined structures solved by X-ray crystallography (light blue) and nuclear magnetic resonance (NMR) spectroscopy (dark blue). To further evaluate the reliability of the structural model, the Ramachandran plot generated by PROCHECK revealed that 93.7% of the amino acid residues fell within the favored regions, 6.3% in allowed regions, and no residues were present in disallowed regions. Collectively, the Z-score and Ramachandran plots indicated that the predicted VirB11 model is of good quality and structurally reliable. We also assessed the sequence homology of VirB11 dimer against the human proteome with E-value ≤ 0.001, and the results revealed no significant homology.

**Fig. 7.**
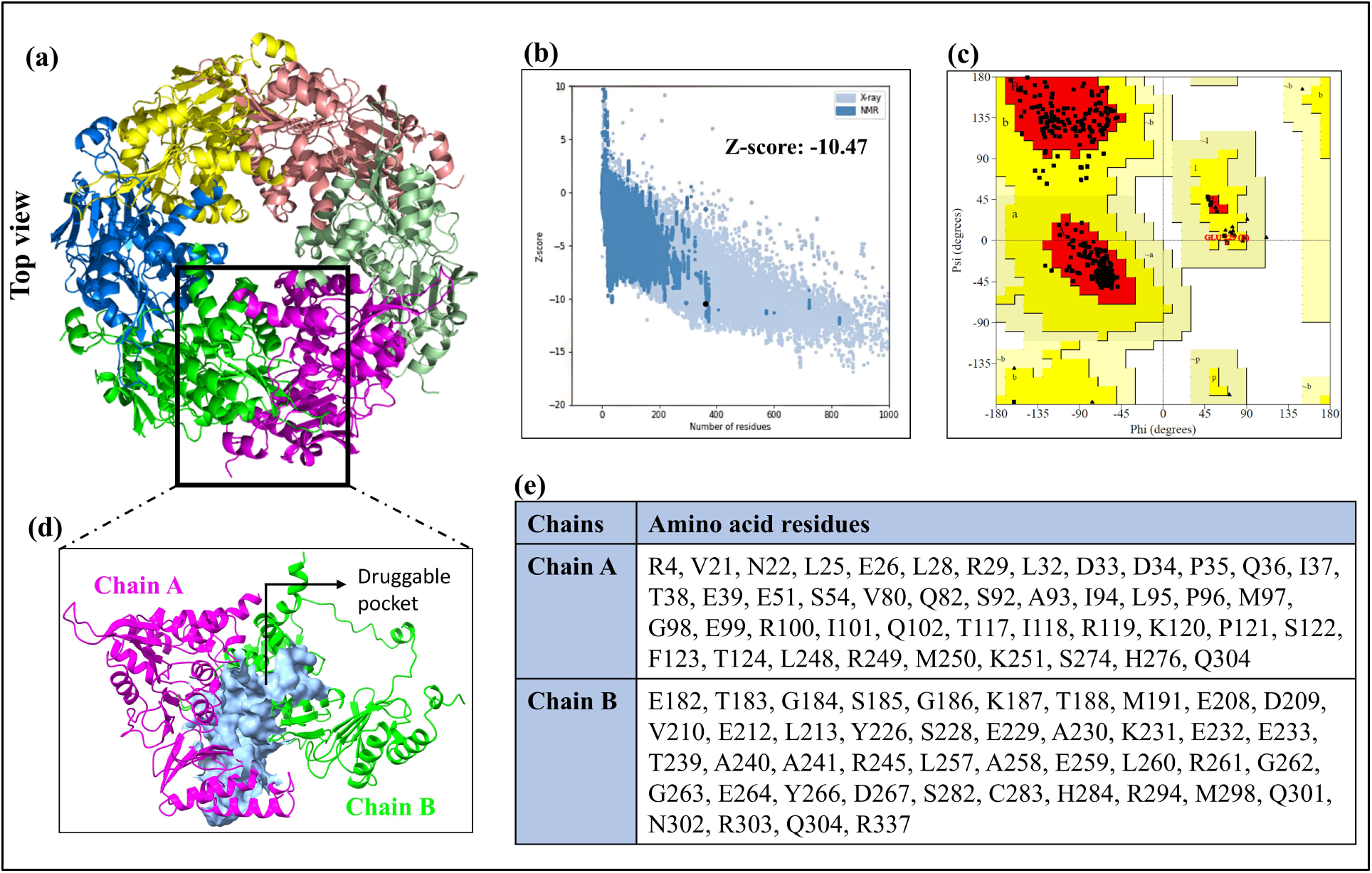
VirB11 model quality assessment and predicted druggable pocket at dimeric interface. **(a)** The hexameric structure of VirB11 modeled using AlphaFold 3 is shown. **(b)** Represents the Z-score plot generated by the ProSA-web server, where the black dot corresponds to the query structure. **(c)** Ramachandran plot generated using the PROCHECK (Red, yellow, and white colors correspond to favored, allowed, and disallowed regions, respectively; squares = non-glycine; and triangles = glycine residues. **(d)** The druggable pocket at VirB11 dimeric interface predicted by DOGSiteScorer is shown. **(e)** The corresponding amino acid residues forming the druggable pocket are displayed.

### 3.8 Prediction of Druggable Pocket at VirB11 dimeric interface

A druggable pocket is a cavity at the monomer–monomer interaction surface that can bind small molecules to disrupt PPI [63]. DOGSiteScorer identified 30 potential druggable pockets within the VirB11 dimeric interface, of which the druggable pocket with the highest druggability score of 0.8 was selected for further analysis. Druggability score close to 1.0 suggests a greater likelihood of the pocket being druggable. The predicted druggable pocket, along with the amino acid residues contributing to its formation, is shown in Figure 7.

### 3.9 Analysis of Molecular Docking Results

Compounds from the DrugBank database were virtually screened against the identified amino acids forming druggable pocket (Figure 8). 130 compounds with docking scores ≤ -7.0 kcal/mol were selected for further analysis. DrugBank is an extensively curated database that integrates in-depth information of drug molecules with their targets and mechanisms of action [64]. Docking scores of ≤ -7.0 kcal/mol are generally indicative of good binding affinity between protein and ligand [56]. These compounds were shortlisted for subsequent ADMET screening.

**Fig. 8.**
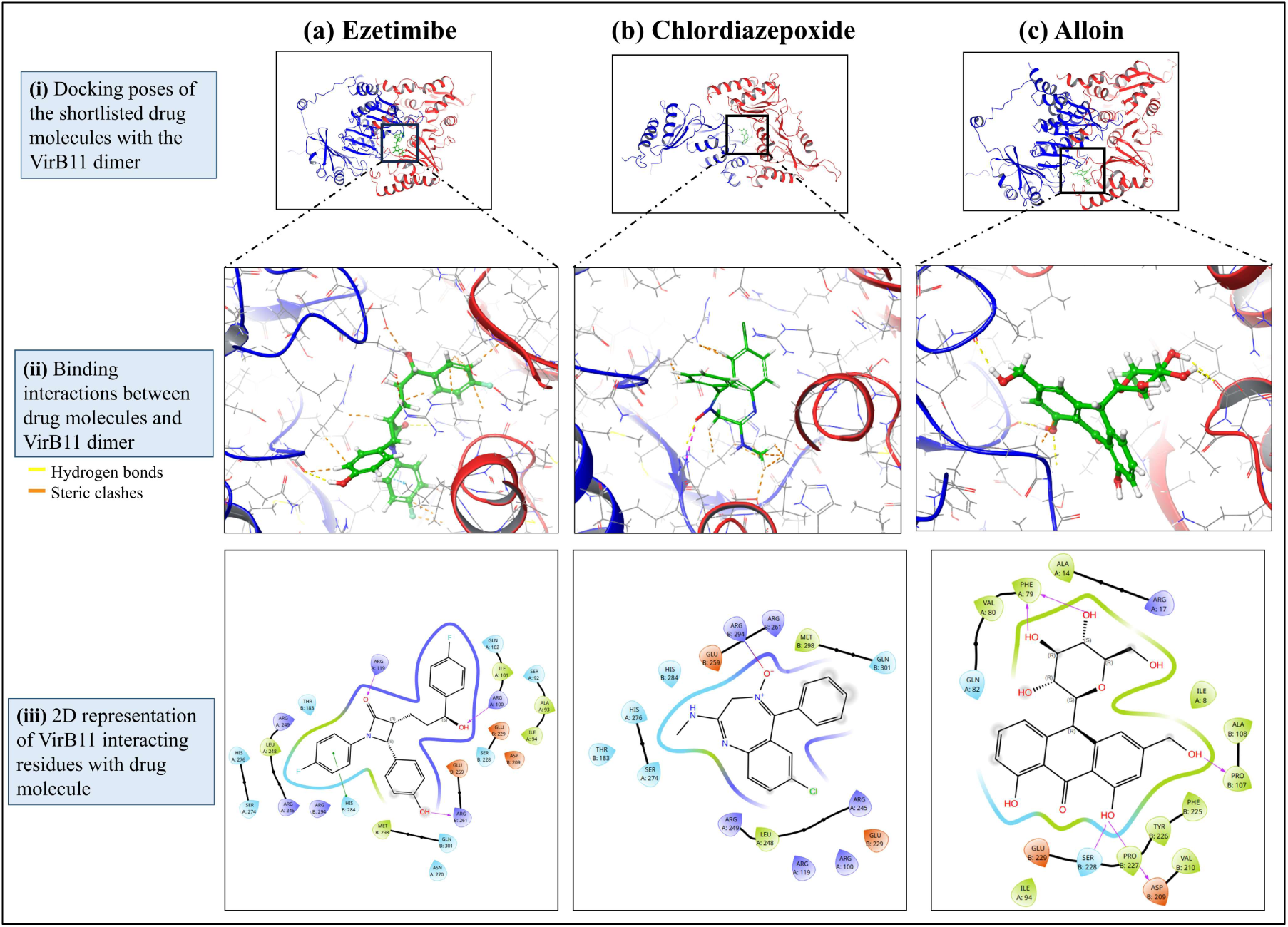
Molecular docking of DrugBank compounds at VirB11 dimeric interface. Molecular docking of DrugBank compounds at the VirB11 dimeric interface. Virtual screening of DrugBank compounds was performed against the predicted druggable pocket, followed by ADMET filtering. **(a)** Ezetimibe, **(b)** Chlordiazepoxide, and **(c)** Alloin docked at the VirB11 dimer interface. For each compound: **(i)** docked pose, **(ii)** enlarged view showing binding interactions (yellow lines, hydrogen bonds; orange lines, steric clashes), and **(iii)** 2D interaction map highlighting ligand–protein contacts are shown.

### 3.10 Prediction of ADMET Properties

A total of 130 compounds targeting the PPI at VirB11 interface, were subjected to ADMET evaluation using the QikProp module of the Schrodinger suite 2025-3. Nine pharmacokinetic criteria were applied for screening according to their corresponding threshold limits. Among the screened compounds, only three compounds– Ezetimibe (DrugBank Id: DB00973), Chlordiazepoxide (DrugBank Id: DB00475), and Alloin (Drug ID: DB15477) – met the ADMET screening criteria (Table 4).

**Table 4:**
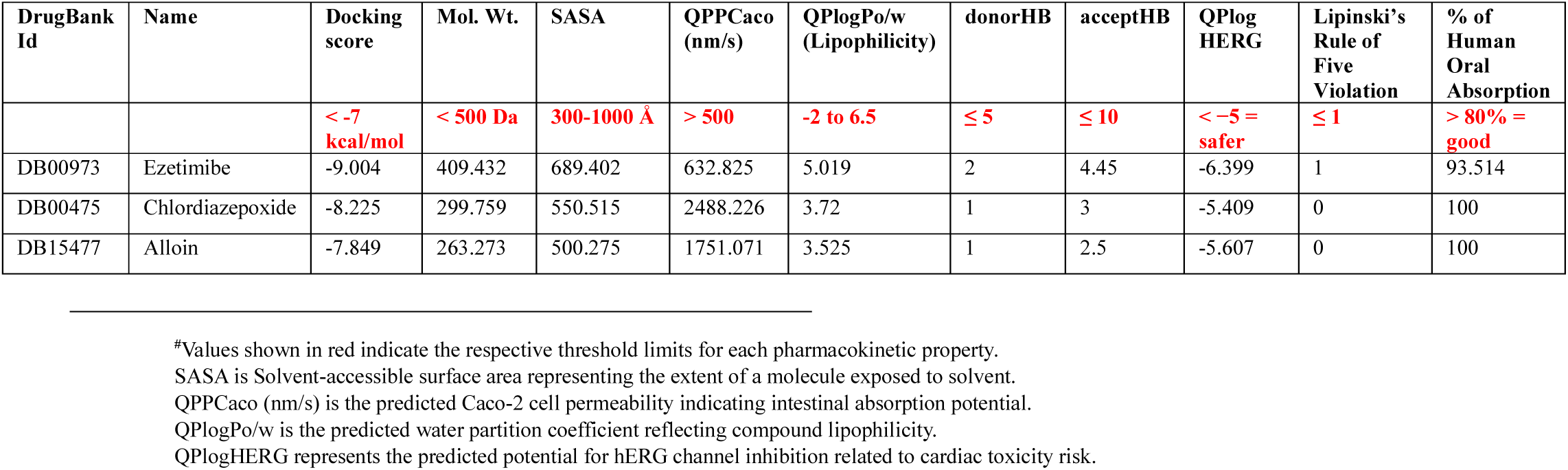
Docking scores and ADMET profile of selected FDA approved compounds from DrugBank after structure-based virtual screening against the VirB11 druggable pocket.

Ezetimibe is an inhibitor of intestinal cholesterol absorption that is used in the treatment of hyperlipidemia [65], Chlordiazepoxide belongs to long-acting benzodiazepines used for the management of mild-to-severe anxiety [66], and Alloin, a stimulant-laxative, is considered to have anti-cancerous properties [67]. All three compounds complied with Lipinski’s rule of five, where Ezetimibe showed one permissible violation, and exhibited docking scores of -9.004, - 8.225, and -7.849, respectively. Chlordiazepoxide and Alloin exhibited 100% human oral absorption, whereas Ezetimibe showed 93.514% (threshold > 80%). Chlordiazepoxide exhibited highest QPPCaco value, suggesting strong intestinal absorption. QPlogPo/w, a measure of lipophilicity, was found to be 5.019 for Ezetimibe and 3.72 for Chlordiazepoxide, and 3.525 for Alloin (threshold range: -2.0 to 6.5), suggesting good membrane permeability. All compounds also showed acceptable QPlogHERG values (ideal value: < -5), indicating a lower risk of hERG channel inhibition (Table 4).

These three compounds were further taken for the calculation of binding energetics and molecular dynamics simulation.

### 3.11 Evaluation of Binding Free Energies of the Docked Complexes

Prime/MM-GBSA analysis depicted that all three shortlisted ligands formed energetically favorable complexes with the VirB11 dimer, with Alloin exhibiting the strongest binding affinity (ΔG_bind_ = -63.20 kcal/mol) (Table 5). Ezetimibe also demonstrated stable binding, with a ΔG_bind_ value of -34.30 kcal/mol, while Chlordiazepoxide showed less negative binding affinity (ΔG_bind_ = -20.51 kcal/mol) among these compounds.

**Table 5:** Prime MM/GBSA-derived binding free energy (ΔG) parameters of selected compounds docked against the VirB11 dimer.

| DrugBank Id | Drug Name | $\Delta G_{\text{bind}}$ | $\Delta G_{\text{cou}}$ | $\Delta G_{\text{cov}}$ | $\Delta G_{\text{lipo}}$ | $\Delta G_{\text{vdw}}$ |
| --- | --- | --- | --- | --- | --- | --- |
| DB00973 | Ezetimibe | -34.30626633 | -21.392 | 13.56488 | -25.4853 | -34.9398 |
| DB00475 | Chlordiazepoxide | -20.51114383 | 18.13062 | 2.983981 | -9.18735 | -35.3765 |
| DB15477 | Alloin | -63.2015885 | -53.6504 | 6.193268 | -15.4131 | -35.9031 |

$\Delta G_{\text{bind}}$ , free energy of binding (kcal/mol); $\Delta G_{\text{cou}}$ , Coulombic energy (kcal/mol); $\Delta G_{\text{cov}}$ , covalent energy (kcal/mol); $\Delta G_{\text{lipo}}$ , hydrophobic energy (kcal/mol); $\Delta G_{\text{vdw}}$ , van der Waals energy (kcal/mol).

Binding affinity of Ezetimibe is largely driven by strong van der Waals and lipophilic interactions, while exhibiting a positive covalent binding. Chlordiazepoxide exhibited weaker overall binding, with unfavorable Coulombic interactions compensated by strong van der Waals forces. Alloin displays the most favorable binding free energy among the selected compounds due to strong Coulombic interactions along with substantial van der Waals contributions, indicating energetically stable association with the VirB11 dimer.

Overall, the MM-GBSA results confirmed that all three ligands formed thermodynamically stable complexes with VirB11 dimer.

### 3.12 Analysis of Molecular Dynamics Simulation

MDS was performed for all three protein-ligand complexes that were selected based on their docking scores (≤ -7.0 kcal/mol) and favorable ADMET properties. For each complex, MDS was performed for 200 ns using the OPLS4 force field at a physiological pH of 7.4 under NPT ensemble conditions with a pressure of 1.01325 bar and a constant temperature of 300K. In the VirB11 dimer – Ezetimibe complex, the protein backbone RMSD began around 2.5 Å and gradually increased to about 4.0 Å within the first 40-50 ns of simulation. Thereafter, the RMSD values remained stable between 3.5 and 4.5 Å for the rest of the simulation, indicating overall structural stability of the protein throughout the simulation. The ligand RMSD showed moderate fluctuations, starting at approximately 3 Å and ranging between 0.6 and 2.5 Å (Figure 9a). These fluctuations reflect conformational adjustments of ligand within the binding pocket without evident dissociation [68]. The RMSF plots revealed that interacting amino acid residues in protein displayed low RMSF values (0.6 to 1.2 Å), indicating that the binding pocket of the VirB11 dimer remains highly stable during the simulation. Notably, key residues (Chain A: R100, R119; and Chain B: S228, E229) involved in hydrogen bonding displayed RMSF values below ∼1.9 Å and remained stable for 36% – 67% of the simulation time. Ligand compactness, assessed by radius of gyration (rGyr; 4.5 Å – 4.95 Å), remained consistent, while SASA values (threshold < 400 Å^2^) ranged from 30 Å^2^ – 95 Å^2^, indicating limited solvent exposure, confirming that Ezetimibe remained deeply embedded within the VirB11 druggable pocket throughout the simulation.

**Fig. 9.**
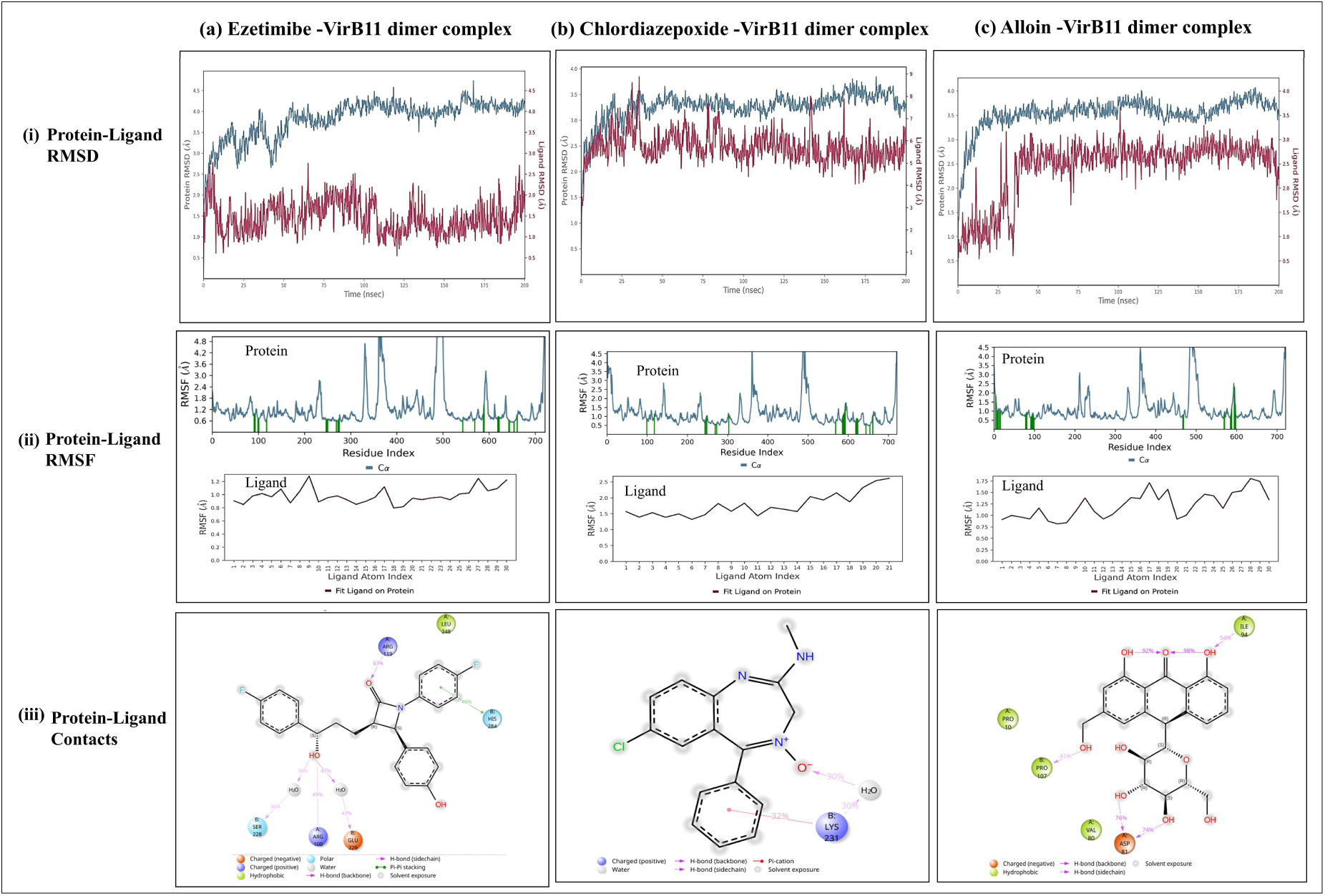
Molecular dynamic simulation of three promising drug candidates for 200 ns. **(a)** Ezetimibe– VirB11 dimer, **(b)** Chlordiazepoxide–VirB11 dimer, and **(c)** Alloin–VirB11 dimer. For each complex, three plots are shown: **(i)** time-dependent RMSD plot showing structural stability, **(ii)** RMSF plot of protein backbone (Cα), where ligand-interacting residues are indicated as green vertical bars, and atom-wise RMSF plot of the ligand indicating stable interactions with the protein, and **(iii)** schematic representation of key protein–ligand interactions persisting for >30% of the simulation time (0–200 ns).

For the VirB11 dimer – Chlordiazepoxide complex, the protein backbone RMSD initially increased to ∼3.5 Å within the first 30-40 ns of the simulation (Figure 9b) during equilibration phase, then gradually stabilized for rest of the simulation with RMSD ranging from 3.0 Å – 3.8 Å. In contrast, the ligand exhibited higher fluctuations compared to the protein, suggesting ligand’s flexible nature. The ligand RMSD began near 3.0 Å and fluctuated between 2.0 and 3.8 Å for the first 50 ns. Thereafter, the RMSD fluctuated between 2.0 Å – 3.5 Å for the remaining simulation, suggesting adaptive conformational changes while remaining bound within the binding pocket. Furthermore, RMSF analysis demonstrated that ligand-interacting amino acid residues exhibited low fluctuation values (0.5 to 1.6 Å), confirming the structural stability of the binding pocket during the simulation. Notably, only K231 (chain B) was involved in forming hydrogen bonds with the ligand for 30% of the simulation time. The ligand’s compactness, evaluated by the rGyr, remained stable in the range of ∼ 3.72 Å – 3.92 Å. SASA values ranged from ∼ 60 Å to 180 Å^2^, indicating that most of the ligand remained deeply buried within the binding pocket with minimal exposure to the solvent.

In the VirB11 dimer – Alloin complex, the protein backbone RMSD increased rapidly from 2.5 Å to 3.5 Å within the first 30 ns, corresponding to the equilibration phase of the system. After this, RMSD values stabilized between 3.0 Å – 4.0 Å, with only minor fluctuations (Figure 9c), indicating overall structural stability of the protein. The ligand RMSD began at around 2.0 Å and hiked to 3.2 Å in first 30 ns, and fluctuated within the range of ∼ 2.0 – 3.5 Å throughout the simulation. RMSF analysis revealed that residues interacting with the ligand exhibited low RMSF values (1.0 Å – 1.5 Å), except N235 (Chain B) with RMSF value of 2.3 Å, supporting the stability of the binding site. D81 and I94 in chain A, and P107 in chain B, are involved in the formation of hydrogen bonds with the ligand. The rGyr, remained stable at approximately 4.15 Å throughout the simulation. SASA analysis showed values ranging from 30 Å² to 120 Å², indicating that a substantial portion of the ligand remained deeply buried within the druggable pocket with minimal solvent exposure.

Taken together, MDS demonstrated that Ezetimibe, Chlordiazepoxide, and Alloin formed stable complexes with the Virb11 dimer over 200 ns, revealing acceptable RMSD and RMSF profiles, stable key intermolecular interactions, ligand compactness, and low solvent exposure (Figure S3). Among the three compounds, Ezetimibe displayed comparatively greater stability, as indicated by lower RMSD and RMSF values, stable interaction profiles, and reduced SASA. Collectively, the dynamic behaviour of the selected compounds supports their potential as promising candidates for further experimental validation and evaluation as repurposed therapeutic agents against *B. melitensis* infection.

## 4. Discussion

Bm is a facultative intracellular pathogen that can survive and replicate within both phagocytic and non-phagocytic host cells. Bm lacks classical virulence determinants such as exotoxins, cytolysins, and exoenzymes. Instead, its pathogenicity primarily relies on key virulence systems: a non-classical LPS that mediates immune evasion, the T4SS that delivers effector proteins, and the BvrR/BvrS two-component system that regulates virulence-associated genes [6]. These systems mediate host-pathogen interaction, and enable the formation of BCVs. BCVs traffic through endosomal compartments, and establish a favourable niche within the endoplasmic reticulum (ER), where bacterial replication occurs [6]. This study structurally characterized one of the key virulence determinants in Bm – the Type IV secretion system (T4SS). Using a computational pipeline, all core VirB proteins were identified, modeled, and assembled into five distinct subcomplexes, leveraging both AlphaFold and template-based approaches. The overall sequence identity between Bm and Ec T4SS proteins was only moderate (∼30-50%), reflecting significant evolutionary divergence at the amino acid level. Despite this, structural alignment with Ec T4SS proteins revealed strong architectural conservation, reflected by low RMSD values (Table 1). Furthermore, the protein models predicted by AlphaFold 3 displayed high confidence, as indicated by reliable pTM and pLDDT scores (Table 1). The stereochemical quality and interface energetics of the modeled subcomplexes were thoroughly validated. PROCHECK analysis showed that nearly all subcomplexes exceeded the 90% favoured region threshold in Ramachandran plots, supporting their geometric reliability. PRODIGY binding energetics calculation revealed strong, stable binding affinities among the interacting protein pairs in Bm T4SS, showing favourable ΔG and K_d_ values comparable to their Ec homologs. Furthermore, structure-based virtual screening identified three potential inhibitors to target PPI at VirB11 interface which is an essential ATPase of Bm T4SS. All three compounds met the screening benchmarks by forming stable, high-affinity complexes with the VirB11 dimer throughout the 200 ns simulations with acceptable RMSD and RMSF profiles, sustained intermolecular interactions, consistent ligand compactness, and limited solvent exposure. Ezetimibe and Alloin showed higher stability compared to Chlordiazepoxide. Ezetimibe showed the highest structural stability during the 200 ns MDS, whereas Alloin displayed the most favorable MM-GBSA binding affinity. All three compounds exhibited favorable pharmacokinetic properties, including high oral bioavailability, metabolic stability, and safety profiles, supporting their potential for repurposing and suggests that they may achieve the intracellular concentrations to inhibit the VirB11 dimerization interface.

Although the overall architecture and assembly energetics of the Bm T4SS closely resemble those reported for Ec, notable structural differences were observed. The VirB3-VirB4 interaction in the IMC of Bm T4SS was substantially stronger than in Ec T4SS as supported by higher number of residue contacts in Bm (53 hydrogen bonds) compared to Ec (24 hydrogen bonds). This potentially indicates adaptations in Bm that either enhance subcomplex stability or alter secretion dynamics. Notably, PDBePISA predicted a positive ΔG value of 0.4 for adjacent VirB4 proteins in IMC. This exception likely reflects a transient interaction, which may be functionally relevant, allowing conformational flexibility necessary for subcomplex assembly or dynamic operation of the T4SS machinery as mentioned in other bacteria [69]. Further, PDBePISA failed to identify residue-level contacts between Ec VirB4 – VirB8 homologs. Interestingly, in Bm, these regions form clear and more defined residue-level contacts, suggesting that Bm T4SS may have evolved a more stabilized VirB4–VirB8 interaction architecture (Table 3). A previous study in Ec T4SS inferred VirB4 – VirB8 contacts using co-evolution analysis, which considers residue–residue pairs with predicted distances ≤12 Å [20]. In context with this, the atomic coordinate file of Ec T4SS showed the distance between the flexible tail regions to be >5 Å as measured in PyMOL 3.0. Thus, PDBePISA could not find any contacts between VirB4 and VirB8 homologs as it could only detect interfaces when atoms fell within a strict distance cutoff (∼ 5 Å), even though the biological interaction exists. The strengthened VirB4 – VirB8 interface in Bm reflects species-specific adaptation of the Bm T4SS according to its intracellular environment. Some interface interactions in Bm showed distinct energetics compared to Ec, suggesting functional or evolutionary divergence within these homologous secretion systems (Tables 2, 3). The Ec T4SS primarily mediates conjugative transfer of plasmid DNA and associated proteins, including relaxases (TraI, TrwC), DNA transfer and replication (Dtr) factors such as TraM, and primases [70]. In contrast, the Bm T4SS functions in both conjugation and virulence by translocating multiple effector proteins into the host cell, thereby reflecting substrate-driven architectural divergence between the two systems [17]. To elucidate the significance of the Bm T4SS, the VirB11 interaction interface was targeted, and the findings validate that disrupting ATPase oligomerization offers an effective anti-virulence strategy to attenuate Bm pathogenicity [29,30]. This study highlights the utility of computational drug repurposing strategies in identifying promising compounds that can interfere with the dimerization of VirB11 which is important for proper functioning of T4SS to export protein effectors. Nevertheless, the study is constrained by the sole use of in silico methodologies – such as homology-based structural prediction, docking analyses, MM-GBSA binding energy estimations, and molecular dynamics simulations – which do not necessarily account for in vivo efficacy, target selectivity, or potential toxicity under physiological conditions.

The identification of thermodynamically stable interactions within Bm T4SS subcomplexes opens several avenues for future research. The interface between several VirB proteins could serve as a novel target for small molecule disruption, with therapeutic implications for brucellosis. The structural differences between Ec and Bm T4SS observed in this study suggest functional adaptations that may impact host-pathogen interactions and secretion efficiency in Bm as compared to other bacteria. The study identified three promising candidates – Ezetimibe, Chlordiazepoxide, and Alloin – to target the assembly of Bm T4SS ATPase that could contribute to the development of anti-virulence therapeutic strategies against Brucellosis. Future experimental studies are required to validate the structures of subcomplexes, evaluate PPI as drug targets, and assess the inhibition of ATPase assembly by the three identified molecules, thereby establishing their relevance in vivo.

## 5. Conclusions

This study structurally characterizes the complete T4SS machinery in Bm, demonstrating high architectural conservation with Ec despite high sequence dissimilarity. Although we generated highly reliable models for predicted subcomplexes, confirmation of these subcomplexes will require experimental structure determination. The identification of stable, functionally relevant protein interfaces within Bm T4SS serve as therapeutic targets to disrupt bacterial virulence. Structure-based virtual screening identified compounds targeting the VirB11 dimerization interface. Three FDA-approved drugs – Ezetimibe, Chlordiazepoxide, and Alloin – showed stable high-affinity binding, with Ezetimibe emerging as the most promising candidate and provides a strong rationale for further experimental validation as a potential anti-virulence agent against Brucellosis.

## Supplementary Materials

Fig. S1. Ramachandran plot analysis of modeled subcomplexes of *B. melitensis* T4SS using PROCHECK. Fig. S2. Orientation of T4SS subcomplexes of *B. melitensis* and *E. coli* within the membrane predicted by the PPM server. Table S1. Identification of T4SS constituent proteins in *B. melitensis* using TXSScan. Table S2. ADMET properties of compounds from DrugBank with docking score < -7 kcal/mol docked against VirB11 druggable pocket.

## Author contributions statement

**Jahnvi Kapoor:** methodology, software, investigation, data analysis, data curation, writing— original draft preparation. **Amisha Panda:** data analysis, visualization, writing—reviewing and editing. **Raman Rajagopal:** methodology, writing—reviewing and editing. **Sanjiv Kumar:** conceptualization, methodology, software, writing—reviewing and editing, supervision. **Anannya Bandyopadhyay**: methodology, data analysis, writing—original draft preparation, writing—reviewing and editing, supervision. All authors reviewed the final draft of the manuscript. All authors contributed to the manuscript revision, read and approved the submitted version.

## Data Availability Statement

The atomic coordinate files (*.pdb) supporting the findings of this study have been deposited in the Zenodo public repository accessible at 10.5281/zenodo.18679107

## Code Availability Statement

The Biopython-based script has been deposited in the Zenodo public repository accessible at 10.5281/zenodo.18679383

## Acknowledgements

JK is a recipient of junior research fellowship from the University Grants Commission (Reference number: 231620067077), Government of India. AP is a recipient of junior research fellowship from the Department of Biotechnology (Grant number: DBT/2023-24/UOD/2326), Government of India.

