## Supplementary material for "Structural Characterization of the Type IV Secretion System in *Brucella melitensis* for Virtual Screening-Based Therapeutic Targeting"

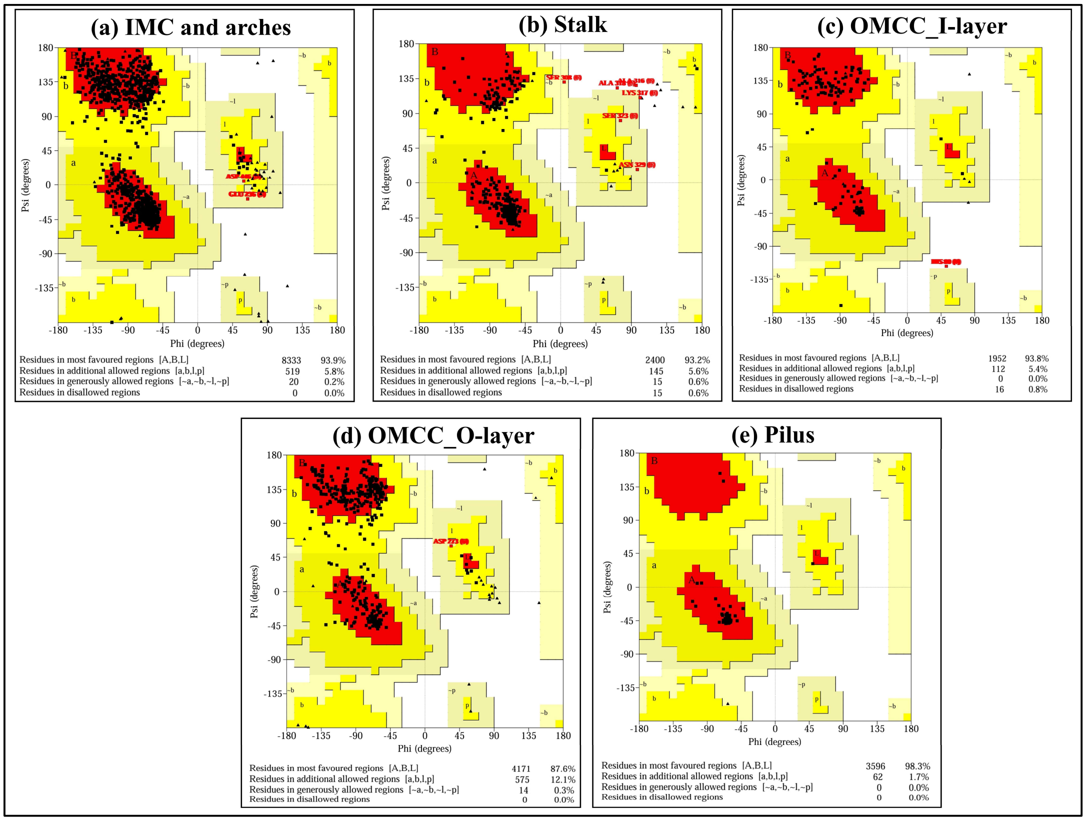


**Fig. S1. Ramachandran plot analysis of modelled subcomplexes of *B. melitensis* T4SS using PROCHECK.** Ramachandran plots represent the backbone dihedral angles (φ and ψ) of amino acid residues for modelled components of the T4SS of *B. melitensis*. The plots show the percentage of residues falling in favoured regions for all the five subcomplexes: **(a)** Inner membrane complex (IMC), **(b)** stalk, **(c)** outer membrane core complex inner layer (OMCC I-layer), **(d)** outer membrane core complex outer layer (OMCC O-layer), and **(e)** pilus.

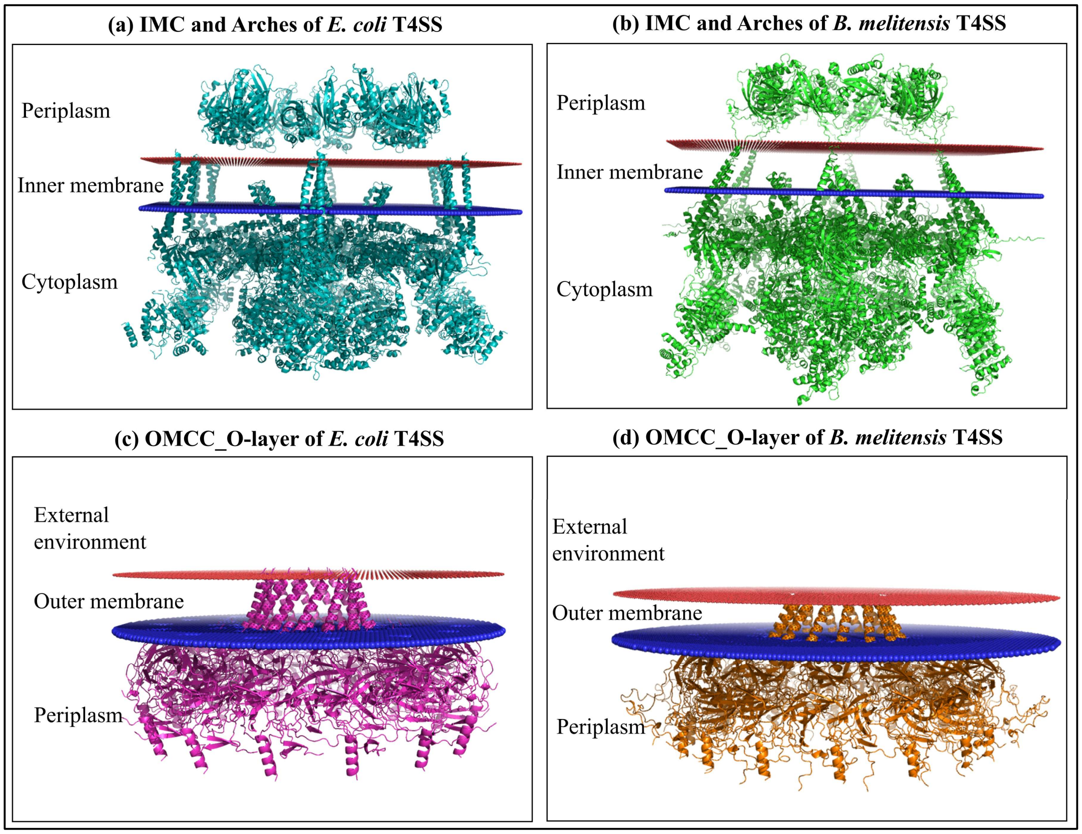


**Fig. S2. Orientation of T4SS subcomplexes of *B. melitensis* and *E. coli* within the membrane predicted by the PPM server.** The membrane insertion and orientation of the IMC and OMCC O-layer were predicted using the PPM server. **(a, b)** The IMC and Arches of *E. coli* and *B. melitensis* show similar overall topology, spanning the inner membrane (IM) with VirB8 tails embedded in the IM and VirB4 anchored to the IM towards periplasmic side. **(c, d)** OMCC O-layer of *E. coli* and *B. melitensis* demonstrate similar orientation patterns. In both the species, helical portion of VirB10 is embedded in the outer membrane (OM) and rest of the domains are towards periplasm.

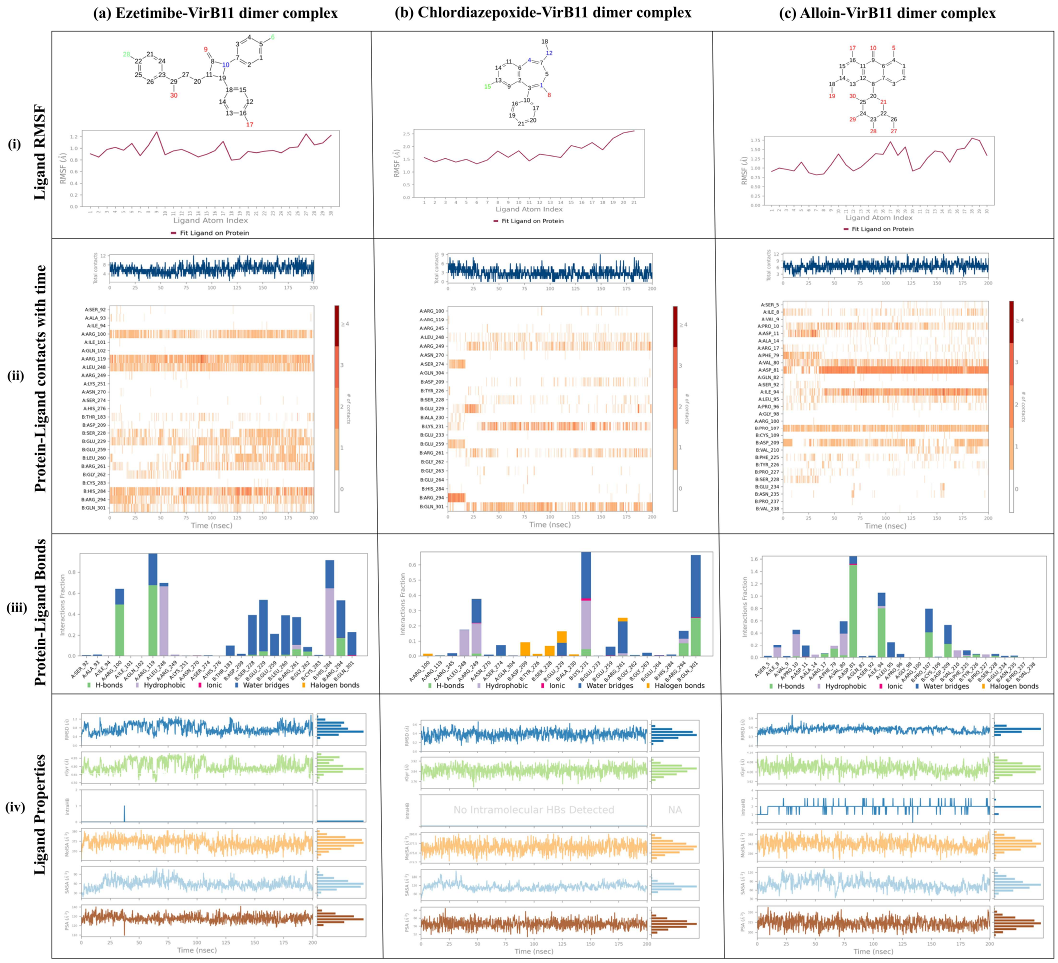


**Fig. S3. Molecular Dynamic Simulation of selected drug candidates with VirB11 dimer.** MDS profile of **(a)** Ezetimibe, **(b)** Chlordiazepoxide, and **(c)** Alloin, with VirB11 dimer are shown. For each complex – **(i)** Ligand RMSF plot with atom-wise flexibility (red = high flexibility, green = high stability, blue = intermediate mobility), **(ii)** A timeline representation of protein-ligand interactions (hydrogen bonds, hydrophobic contacts, ionic interactions, water bridges, and halogen bonds) is shown, **(iii)** Protein-ligand interaction profile throughout the simulation, and **(iv)** Dynamics of ligand's physicochemical properties during the simulation period (0-200 ns) is shown.

**Table S1.** Identification of T4SS constituent proteins in *B. melitensis* using TXSScan

| Hit Id | Gene name | Hit E-value | Hit position | Profile coverage |
|---|---|---|---|---|
| **WP_004680997.1** | T4SS_T_virB1 | 7.30E-06 | 1363 | 0.811 |
| **WP_004681227.1** | T4SS_T_virB1 | 1.30E-66 | 1412 | 1 |
| **WP_002967165.1** | T4SS_T_virB2 | 1.60E-28 | 1031 | 0.894 |
| **WP_002966512.1** | T4SS_T_virB3 | 3.20E-22 | 931 | 0.877 |
| **WP_004681223.1** | T4SS_T_virb4 | 1.10E-225 | 1411 | 0.828 |
| **WP_002966514.1** | T4SS_T_virB5 | 2.90E-55 | 932 | 1 |
| **WP_004686828.1** | T4SS_T_virB6 | 3.40E-35 | 2706 | 0.746 |
| **WP_002966517.1** | T4SS_T_virB8 | 3.30E-72 | 934 | 0.961 |
| **WP_002966518.1** | T4SS_T_virB9 | 1.00E-79 | 935 | 0.974 |
| **WP_004685137.1** | T4SS_T_virB10 | 5.80E-33 | 2580 | 0.664 |

**Table S2.** ADMET properties of compounds from DrugBank with docking score < -7 kcal/mol docked against VirB11 druggable pocket

| DrugBank Id | Name | Docking score | mol_MW | SASA | QPPCaco (nm/s) | QPlogPo/w (Lipophilicity) | donorHB | acceptHB | QPlogHERG | Lipinski's Rule of Five Violation | % of Human Oral Absorption |
|---|---|---|---|---|---|---|---|---|---|---|---|
| DB16019 | Gallium Ga-68 gozetotide | -12.56 | 947.004 | 1565.012 | 0 | -0.842 | 10.5 | 23 | 5.046 | 3 | 0 |
| DB00157 | NADH | -11.33 | 665.446 | 975.294 | 0.004 | -3.285 | 8 | 25.2 | -3.444 | 3 | 0 |
| DB06636 | Isavuconazonium | -10.69 | – | – | – | – | – | – | – | – | – |
| DB11183 | Light green SF yellowish | -10.02 | – | – | – | – | – | – | – | – | – |
| DB06796 | Mangafodipir | -9.955 | 638.461 | 944.859 | 0 | -2.774 | 8 | 21.5 | 3.389 | 3 | 0 |
| DB06441 | Cangrelor | -9.827 | 776.353 | 953.327 | 0.047 | 2.088 | 3 | 20.6 | 1.444 | 3 | 0 |
| DB14879 | Cefiderocol | -9.518 | – | – | – | – | – | – | – | – | – |
| DB01328 | Cefonicid | -9.439 | 542.556 | 796.404 | 0.08 | 0.097 | 3.25 | 14.95 | -1.151 | 2 | 0 |
| DB01601 | Lopinavir | -9.423 | 628.81 | 1089.182 | 674.821 | 6.07 | 4 | 9.45 | -5.65 | 2 | 87.208 |
| DB03310 | Glutathione disulfide | -9.218 | 612.626 | 989.819 | 0 | -6 | 9 | 18 | 5.139 | 3 | 0 |
| DB03147 | Flavin adenine dinucleotide | -9.008 | 785.557 | 1068.125 | 0.002 | -3.016 | 8 | 27.7 | -3.235 | 3 | 0 |
| DB00973 | Ezetimibe | -9.004 | 409.432 | 689.402 | 632.825 | 5.019 | 2 | 4.45 | -6.399 | 1 | 93.514 |
| DB04711 | Iodipamide | -8.971 | 1139.767 | 859.366 | 3.373 | 5.755 | 4 | 9 | -2.398 | 2 | 44.174 |
| DB00743 | Gadobenic acid | -8.948 | 513.5 | 834.117 | 0 | -2.339 | 5 | 17.7 | 0.636 | 2 | 0 |
| DB00563 | Methotrexate | -8.761 | 454.444 | 742.537 | 0.045 | -1.78 | 6.25 | 11.75 | -2.11 | 2 | 0 |
| DB13074 | Macimorelin | -8.76 | 474.561 | 812.79 | 6.937 | 2.224 | 4.25 | 5.75 | -3.53 | 1 | 42.065 |
| DB01329 | Cefoperazone | -8.579 | 645.664 | 979.551 | 0.093 | 0.577 | 2.25 | 15.5 | -2.016 | 2 | 0 |
| DB15617 | Ferric derisomaltose | -8.545 | 506.457 | 798.21 | 0.327 | -6.17 | 12 | 27.2 | -5.195 | 3 | 0 |
| DB06590 | Ceftaroline fosamil | -8.523 | – | – | – | – | – | – | – | – | – |
| DB00605 | Sulindac | -8.516 | 356.411 | 622.173 | 4.97 | 3.421 | 1 | 6 | -3.627 | 0 | 59.438 |
| DB00183 | Pentagastrin | -8.507 | 767.896 | 1220.122 | 0.553 | 3.34 | 4.75 | 13.25 | 0.499 | 3 | 3.029 |
| DB05521 | Telaprevir | -8.506 | 679.858 | 1152.962 | 108.616 | 3.818 | 0.75 | 14.25 | -2.78 | 2 | 59.824 |
| DB06705 | Gadofosveset trisodium | -8.478 | 737.696 | 987.09 | 0 | -0.137 | 6 | 21 | 2.063 | 3 | 0 |
| DB00689 | Cephaloglycin | -8.475 | 405.425 | 678.413 | 1.455 | -1.174 | 3.25 | 9.25 | -3.048 | 0 | 22.983 |
| DB12615 | Plazomicin | -8.393 | 592.688 | 881.391 | 0.194 | -5.636 | 14 | 21.9 | -7.492 | 3 | 0 |
| DB06813 | Pralatrexate | -8.333 | 477.479 | 823.459 | 0.065 | 1.412 | 6.75 | 10.75 | -2.977 | 2 | 0 |

| | | | | | | | | | | | |
|---|---|---|---|---|---|---|---|---|---|---|---|
| DB09425 | Indium In-111 pentetate | -8.319 | 393.35 | 649.158 | 0 | -4.266 | 5 | 16 | 2.461 | 1 | 0 |
| DB14075 | Imidurea | -8.317 | 388.296 | 587.7 | 0.433 | -2.215 | 2.5 | 8.9 | -1.387 | 2 | 0 |
| DB00471 | Montelukast | -8.287 | 586.187 | 1023.465 | 130.854 | 9.526 | 2 | 4.25 | -6.51 | 2 | 94.691 |
| DB09313 | Ioxaglic acid | -8.268 | 1268.886 | 988.73 | 3.276 | 3.211 | 5.25 | 15.95 | -4.223 | 3 | 16.093 |
| DB11979 | Elagolix | -8.255 | 631.598 | 917.915 | 48.28 | 5.079 | 2 | 7.75 | -6.186 | 2 | 60.906 |
| DB00475 | Chlordiazepoxide | -8.225 | 299.759 | 550.515 | 2488.226 | 3.72 | 1 | 3 | -5.409 | 0 | 100 |
| DB02772 | Sucrose | -8.146 | 342.299 | 528.582 | 11.688 | -3.698 | 8 | 16.8 | -3.434 | 2 | 0 |
| DB11217 | Arbutin | -8.017 | 272.254 | 490.771 | 71.832 | -0.99 | 5 | 10 | -4.315 | 0 | 54.371 |
| DB09038 | Empagliflozin | -7.997 | 450.915 | 724.618 | 276.014 | 1.967 | 4 | 10.95 | -5.674 | 0 | 82.15 |
| DB09332 | Kappadione | -7.97 | 334.159 | 501.648 | 0.523 | 1.115 | 4 | 10 | 2.914 | 0 | 28.435 |
| DB05039 | Indacaterol | -7.962 | 392.497 | 698.085 | 64.909 | 2.592 | 4 | 6.45 | -6.106 | 0 | 74.558 |
| DB01200 | Bromocriptine | -7.904 | 654.602 | 887.711 | 126.195 | 3.406 | 2.25 | 10.75 | -3.748 | 1 | 71.532 |
| DB00293 | Raltitrexed | -7.887 | 458.488 | 788.963 | 0.416 | 2.406 | 3.25 | 10.25 | -2.669 | 0 | 34.22 |
| DB14703 | Dexamethasone metasulfobenzoate | -7.879 | 576.633 | 831.118 | 3.192 | 2.534 | 3 | 12.45 | -4.104 | 1 | 37.846 |
| DB00776 | Oxcarbazepine | -7.877 | 252.272 | 462.974 | 322.641 | 1.264 | 2 | 4 | -3.31 | 0 | 79.25 |
| DB00966 | Telmisartan | -7.858 | 514.626 | 887.236 | 227.343 | 7.787 | 1 | 5 | -5.901 | 2 | 88.806 |
| DB14185 | Aripiprazole lauroxil | -7.857 | 660.723 | 1240.246 | 379.99 | 9.003 | 0 | 8.75 | -8.715 | 2 | 100 |
| DB15477 | Alloin | -7.849 | 418.399 | 629.721 | 11.821 | -0.406 | 5 | 11.7 | -4.785 | 1 | 30.808 |
| DB14914 | Flortaucipir F-18 | -7.849 | 263.273 | 500.275 | 1751.071 | 3.525 | 1 | 2.5 | -5.607 | 0 | 100 |
| DB08934 | Sofosbuvir | -7.846 | 529.458 | 807.979 | 75.806 | 1.211 | 3 | 14.9 | -6.164 | 2 | 41.764 |
| DB09065 | Cobicistat | -7.818 | 776.023 | 1209.811 | 95.239 | 6.535 | 2.25 | 12.95 | -6.379 | 3 | 61.753 |
| DB16216 | Lazertinib | -7.798 | 554.65 | 917.93 | 323.544 | 4.324 | 2 | 11.45 | -8 | 2 | 71.269 |
| DB11190 | Pantethine | -7.778 | 554.716 | 930.441 | 8.704 | -1.19 | 6 | 15.8 | -0.181 | 3 | 0 |
| DB00385 | Valrubicin | -7.736 | 723.653 | 1030.153 | 29.057 | 2.542 | 3 | 16.6 | -5.144 | 2 | 42.101 |
| DB14575 | Eslicarbazepine | -7.712 | 254.288 | 468.972 | 340.055 | 1.326 | 3 | 3.7 | -3.212 | 0 | 80.018 |
| DB06210 | Eltrombopag | -7.708 | 442.473 | 792.841 | 35.561 | 4.441 | 2 | 7.25 | -5.277 | 0 | 80.71 |
| DB12010 | Fostamatinib | -7.678 | 580.465 | 907.61 | 6.524 | 2.813 | 4 | 15 | -3.26 | 2 | 32.079 |
| DB00598 | Labetalol | -7.672 | 328.41 | 603.077 | 39.717 | 2.71 | 4 | 5.45 | -5.954 | 0 | 71.433 |
| DB09119 | Eslicarbazepine acetate | -7.662 | 296.325 | 542.561 | 317.04 | 2.15 | 2 | 4 | -3.86 | 0 | 84.302 |

| | | | | | | | | | | | |
|---|---|---|---|---|---|---|---|---|---|---|---|
| DB00118 | Ademetionine | -7.646 | – | – | – | – | – | – | – | – | – |
| DB11596 | Levoleucovorin | -7.612 | 473.444 | 784.298 | 0.018 | -0.768 | 7.25 | 14.25 | -2.456 | – | 0 |
| DB19360 | Latanoprost acid | -7.593 | 390.519 | 752.497 | 24.925 | 3.616 | 4 | 7.1 | -4.148 | – | 73.116 |
| DB01112 | Cefuroxime | -7.569 | 424.384 | 680.876 | 4.477 | 0.105 | 3.25 | 11.95 | -3.747 | – | 26.254 |
| DB00158 | Folic acid | -7.545 | 441.402 | 731.339 | 0.034 | -0.415 | 6.25 | 12.75 | -2.44 | – | 0 |
| DB00188 | Bortezomib | -7.515 | – | – | – | – | – | – | – | – | – |
| DB03247 | Flavin mononucleotide | -7.511 | 456.348 | 657.12 | 0.255 | -1.008 | 6 | 15.6 | -0.936 | – | 0 |
| DB09049 | Naloxegol | -7.491 | 651.793 | 1028.465 | 819 | 3.23 | 2 | 17.85 | -7.07 | – | 72.083 |
| DB11327 | Dipyrithione | -7.488 | 252.305 | 460.964 | 2005.771 | 3.385 | 0 | 2 | -5.139 | – | 100 |
| DB01331 | Cefoxitin | -7.484 | 427.446 | 643.854 | 4.147 | 0.42 | 3.25 | 9.5 | -1.587 | – | 40.459 |
| DB00390 | Digoxin | -7.481 | 780.948 | 1037.385 | 38.497 | 1.027 | 6 | 22.45 | -5.347 | – | 22.459 |
| DB15982 | Berotralstat | -7.478 | 562.569 | 933.788 | 15.296 | 4.959 | 4 | 7.5 | -9.08 | – | 64.224 |
| DB00630 | Alendronic acid | -7.467 | 249.097 | 405.236 | 0.046 | -2.031 | 3 | 7.75 | 3.549 | – | 0 |
| DB00650 | Leucovorin | -7.46 | 473.444 | 756.783 | 0.022 | -0.762 | 7.25 | 14.25 | -2 | – | 0 |
| DB00399 | Zoledronic acid | -7.456 | 272.091 | 408.853 | 0.395 | 0.501 | 1 | 8.75 | 3.98 | – | 22.669 |
| DB04868 | Nilotinib | -7.443 | 529.523 | 896.842 | 671.75 | 5.898 | 2 | 8 | -8.133 | – | 86.165 |
| DB01102 | Arbutamine | -7.435 | 317.384 | 638.466 | 29.982 | 1.503 | 5 | 5.45 | -6.628 | – | 62.179 |
| DB01632 | 5-O-phosphono-alpha-D-ribofuranosyl diphosphate | -7.411 | 390.071 | 547.864 | 0.01 | -1.871 | 4 | 17.1 | 4.903 | – | 0 |
| DB09570 | Ixazomib | -7.403 | – | – | – | – | – | – | – | – | – |
| DB05273 | Samarium (153Sm) lexidronam | -7.4 | 436.125 | 614.864 | 0 | -4.965 | 8 | 24 | 8.838 | – | 0 |
| DB01204 | Mitoxantrone | -7.388 | 444.486 | 791.956 | 0.751 | 0.503 | 4 | 9.9 | -7.625 | – | 14.713 |
| DB04861 | Nebivolol | -7.388 | 405.441 | 708.431 | 538.459 | 3.806 | 3 | 6.4 | -6.897 | – | 100 |
| DB00274 | Cefmetazole | -7.383 | 471.523 | 723.854 | 3.794 | 0.666 | 1.25 | 12 | -1.797 | – | 28.252 |
| DB12362 | Diaminopropanol tetraacetic acid | -7.383 | 322.271 | 519.428 | 0.001 | -3.962 | 5 | 13.7 | 2.322 | – | 0 |
| DB09280 | Lumacaftor | -7.368 | 452.413 | 722.174 | 87.576 | 4.276 | 2 | 7.5 | -4.521 | – | 86.746 |
| DB00642 | Pemetrexed | -7.366 | 427.416 | 717.347 | 0.087 | 0.859 | 6.25 | 9.75 | -2.163 | – | 0 |
| DB00131 | Adenosine phosphate | -7.362 | 347.224 | 529.974 | 0.365 | -1.659 | 6 | 14.1 | -0.664 | – | 0 |
| DB08834 | Tauroursodeoxycholic acid | -7.356 | 499.705 | 786.391 | 9.513 | 2.949 | 3 | 8.9 | -1.47 | – | 61.724 |

| | | | | | | | | | | | |
|---|---|---|---|---|---|---|---|---|---|---|---|
| DB06817 | Raltegravir | -7.353 | 444.421 | 749.659 | 80.064 | 1.753 | 2 | 11.25 | -5.928 | – | 58.317 |
| DB16703 | Belumosudil | -7.348 | 452.515 | 773.449 | 399.845 | 4.023 | 3 | 6.75 | -5.801 | – | 100 |
| DB14805 | Piflufolastat F 18 | -7.328 | 442.4 | 778.395 | 0.014 | 1.499 | 4.5 | 10 | 0.726 | – | 0 |
| DB00535 | Cefdinir | -7.322 | 395.407 | 618.604 | 2.735 | 0.393 | 3.25 | 9.95 | -3.098 | – | 37.064 |
| DB00342 | Terfenadine | -7.319 | 471.681 | 862.545 | 547.316 | 6.802 | 2 | 4.45 | -7.95 | – | 100 |
| DB08816 | Ticagrelor | -7.312 | 522.569 | 883.205 | 141.496 | 3.247 | 4 | 11.3 | -6.441 | – | 71.491 |
| DB14761 | Remdesivir | -7.298 | 602.583 | 999.442 | 14.466 | 1.536 | 5 | 16.65 | -7.67 | – | 30.791 |
| DB09268 | Picosulfuric acid | -7.298 | 437.438 | 673.972 | 0.688 | 1.823 | 2 | 10 | -2.699 | – | 34.712 |
| DB01133 | Tiludronic acid | -7.288 | 318.605 | 471.988 | 1.21 | 2.352 | 0 | 6.5 | 3.082 | – | 42.203 |
| DB09090 | Pinaverium | -7.287 | – | – | – | – | – | – | – | – | – |
| DB01026 | Ketoconazole | -7.282 | 531.438 | 831.756 | 867.511 | 4.381 | 0 | 8.25 | -5.355 | – | 92.23 |
| DB04209 | Dequalinium | -7.279 | 456.673 | 891.309 | 941.838 | 7.377 | 3 | 3 | -7.484 | – | 100 |
| DB09076 | Umeclidinium | -7.268 | – | – | – | – | – | – | – | – | – |
| DB04703 | Hesperidin | -7.263 | 610.568 | 904.749 | 4.796 | -1.247 | 7 | 20.05 | -6.335 | – | 0 |
| DB09267 | Strontium ranelate | -7.252 | 342.28 | 530.339 | 0.024 | 0.149 | 4 | 10.5 | 3.486 | – | 0 |
| DB00314 | Capreomycin | -7.252 | 668.712 | 954.602 | 0 | -5.26 | 10 | 14.7 | -0.813 | – | 0 |
| DB13345 | Dihydroergocristine | -7.236 | 611.739 | 902.461 | 139.993 | 3.66 | 2.25 | 10.75 | -4.66 | – | 73.828 |
| DB13680 | Naftazone | -7.236 | 215.211 | 442.975 | 56.952 | 0.741 | 2 | 2.5 | -3.439 | – | 62.706 |
| DB14650 | Menadiol diphosphate | -7.209 | 334.159 | 488.585 | 0.567 | 1.097 | 4 | 10 | 3.205 | – | 28.952 |
| DB00923 | Ceforanide | -7.206 | 519.549 | 791.721 | 0.031 | -1.363 | 4.25 | 12.25 | -1.511 | – | 0 |
| DB08882 | Linagliptin | -7.202 | 472.549 | 798.234 | 188.579 | 3.715 | 2 | 8.5 | -6.737 | – | 89.423 |
| DB11611 | Lifitegrast | -7.198 | 615.484 | 883.319 | 20.006 | 4.681 | 1.25 | 11.25 | -5.224 | – | 64.685 |
| DB07565 | Chloramphenicol succinate | -7.197 | 423.206 | 614.168 | 2.441 | 0.721 | 3 | 9.2 | -1.267 | – | 38.101 |
| DB08976 | Floctafenine | -7.195 | 406.361 | 681.697 | 420.867 | 4.055 | 2 | 5.9 | -6.283 | – | 100 |
| DB00779 | Nalidixic acid | -7.189 | 232.238 | 462.559 | 96.563 | 1.59 | 0 | 4.5 | -2.213 | – | 71.781 |
| DB01333 | Cefradine | -7.18 | 349.404 | 609.623 | 3.533 | -1.003 | 3.25 | 7.25 | -2.34 | – | 30.88 |
| DB00503 | Ritonavir | -7.172 | 720.943 | 1148.087 | 358.89 | 6.818 | 3.25 | 10.95 | -5.653 | – | 73.718 |
| DB00567 | Cephalexin | -7.155 | 347.388 | 593.79 | 3.437 | -1.185 | 3.25 | 7.25 | -2.585 | – | 29.603 |
| DB00282 | Pamidronic acid | -7.15 | 235.07 | 374.406 | 0.037 | -2.362 | 3 | 7.75 | 3.795 | – | 0 |
| DB08909 | Glycerol phenylbutyrate | -7.147 | 530.66 | 1085.06 | 526.616 | 8.5 | 0 | 6 | -9.437 | – | 100 |

| | | | | | | | | | | | |
|---|---|---|---|---|---|---|---|---|---|---|---|
| DB00456 | Cefalotin | -7.144 | 396.432 | 668.076 | 11.011 | 1.931 | 1.25 | 8.25 | -2.176 | – | 56.901 |
| DB00303 | Ertapenem | -7.141 | 475.515 | 772.249 | 0.089 | -0.486 | 4 | 11.2 | -2.856 | – | 5.268 |
| DB06374 | Elacestrant | -7.126 | 458.642 | 830.84 | 605.144 | 6.503 | 2 | 4 | -7.212 | – | 100 |
| DB01609 | Deferasirox | -7.122 | 373.367 | 647.842 | 22.136 | 3.138 | 3 | 6 | -4.786 | – | 69.396 |
| DB01003 | Cromoglicic acid | -7.12 | 468.373 | 719.405 | 0.417 | 1.897 | 3 | 12.2 | -2.701 | – | 18.303 |
| DB01987 | Cocarboxylase | -7.118 | | | | | | | | – | |
| DB00452 | Framycetin | -7.112 | 614.649 | 890.569 | 0.001 | -9.996 | 19 | 28.1 | -9.645 | – | 0 |
| DB05667 | Levoketoconazole | -7.108 | 531.438 | 851.625 | 895.608 | 4.361 | 0 | 8.25 | -5.639 | – | 92.359 |
| DB09134 | Ioversol | -7.103 | 807.116 | 790.605 | 4.534 | -1.524 | 8 | 18.2 | -5.06 | – | 0 |
| DB00878 | Chlorhexidine | -7.091 | 505.452 | 929.801 | 23.153 | 3.419 | 10 | 7 | -7.632 | – | 45.469 |
| DB16907 | Zoledronate D,L-Lysine Monohydrate | -7.085 | 272.091 | 409.099 | 0.396 | 0.469 | 1 | 8.75 | 3.917 | – | 22.491 |
| DB00950 | Fexofenadine | -7.076 | 501.664 | 885.074 | 23.195 | 4.079 | 3 | 6.45 | -6.334 | – | 62.31 |
| DB00770 | Alprostadil | -7.073 | 354.486 | 736.606 | 19.316 | 2.968 | 3 | 7.4 | -3.235 | – | 67.337 |
| DB00496 | Darifenacin | -7.03 | 426.557 | 763.848 | 282.588 | 4.55 | 2 | 5.25 | -6.43 | – | 100 |
| DB00735 | Naftifine | -7.019 | 287.404 | 604.352 | 2470.586 | 5.596 | 0 | 2 | -7.285 | – | 100 |
| DB04570 | Latamoxef | -7.015 | 520.473 | 795.481 | 0.153 | -0.426 | 2 | 12.7 | -0.986 | – | 0 |
| DB00701 | Amprenavir | -7.01 | 505.628 | 826.394 | 317.283 | 3.193 | 3.5 | 11.4 | -6.422 | – | 77.453 |

Compounds shortlisted after screening are highlighted in yellow.
– These values were not predicted by QikProp module for the given compounds.
SASA is Solvent-accessible surface area representing the extent of a molecule exposed to solvent.
QPPCaco (nm/s) is the predicted Caco-2 cell permeability indicating intestinal absorption potential.
QPlogPo/w is the predicted water partition coefficient reflecting compound lipophilicity.
QPlogHERG represents the predicted potential for hERG channel inhibition related to cardiac toxicity risk.
